# UVB modifies skin immune-stroma cross-talk and promotes effector T cell recruitment during cryptic *Leishmania donovani* infection

**DOI:** 10.1101/2023.02.03.526940

**Authors:** Marcela Montes de Oca, Shoumit Dey, Katrien Van Bocxlaer, Helen Ashwin, Najmeeyah Brown, Elmarie Myburgh, Nidhi S Dey, Gulab Fatima Rani, Edward Muscutt, Mohamed Osman, Damian Perez-Mazliah, Sally James, Lesley Gilbert, Mitali Chatterjee, Paul M Kaye

## Abstract

Many parasites of significant public health importance assume skin residency without causing overt pathlogy. How immune and stromal cells respond to such “cryptic” infections and how exposure to UVB alters such responses in poorly understood. We combined scRNA-seq, spatial transcriptomics and inferential network analysis to address these questions in a model of cryptic skin infection by *Leishmania donovani*. In infected C57BL/6 mice, p-selectin and CXCL12 interactions dominate intercellular communication between leucocytes, fibroblast and endothelial cells, but effector T cell function remains muted. Following UVB exposure, increased numbers of IFNγ^+^ CD4^+^ Th1 cells and NK cells enter the skin, communicating with stromal cells via CCL5-CCR5 and LFA-1-ICAM1/2. However, spatial mapping indicated that Th1 cells and macrophages occupied distinct niches after UVB exposure, likely limiting effector function. Our data provide the first holistic view of the immune landscape during cryptic *L. donovani* infection and demonstrate how UVB exposure fundamentally reshapes this response.

**GRAPHICAL ABSTRACT:** 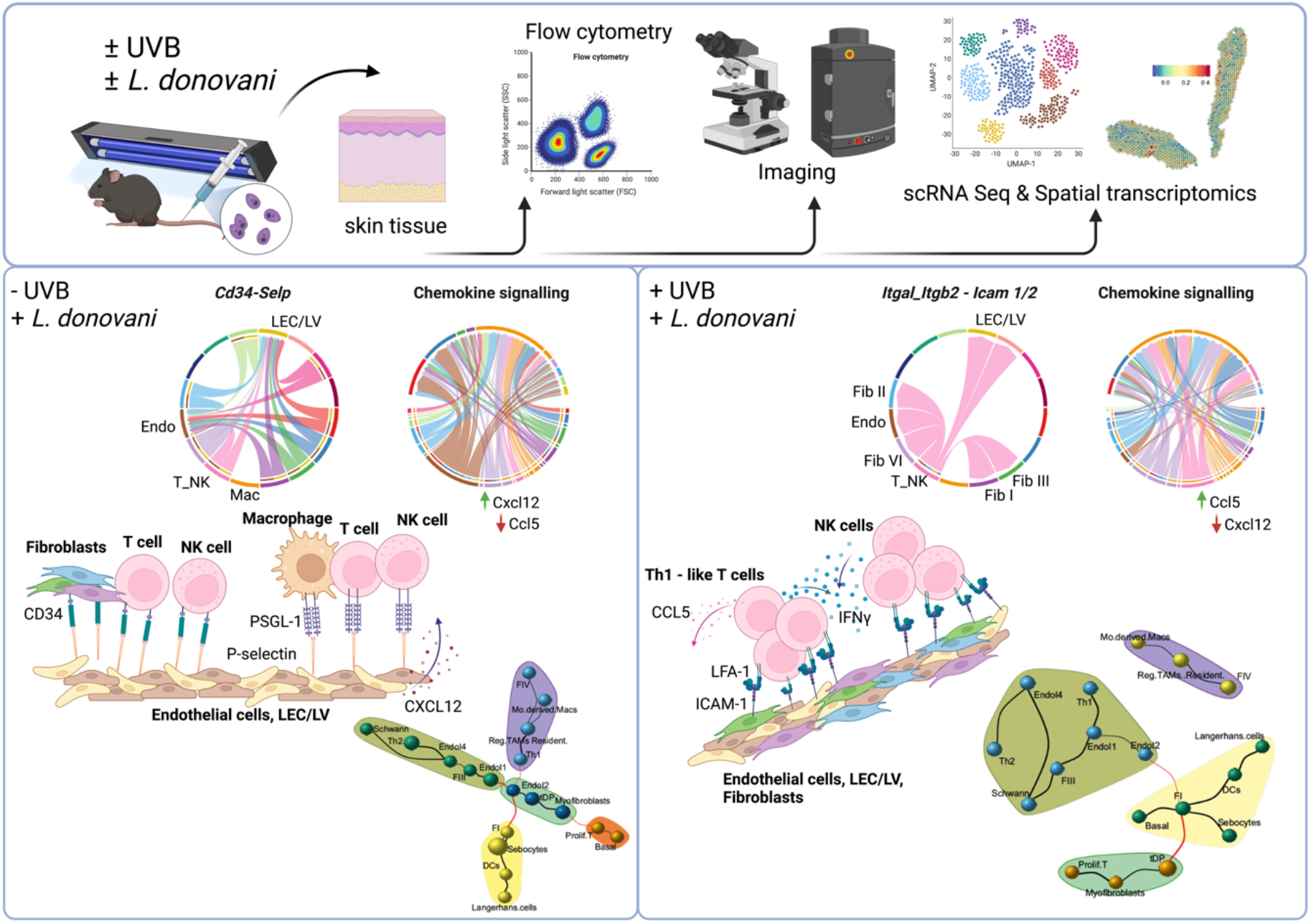

## INTRODUCTION

Trypanosomatid parasites cause a range of globally important diseases, including leishmaniasis, African trypanosomiasis and Chagas disease, with significant consequences for health and economic prosperity. Whilst more notable for causing systemic disease, recent studies in mice and humans have highlighted the skin as an important site of parasitism by many trypanosomatids^1–4^ including *Leishmania donovani* the causative agent of visceral leishmaniasis ^5–9^. Such skin infections generally proceed in the absence of overt pathology yet may facilitate parasite transmission and provide a challenge for chemotherapy ^10^. Few studies, however, have systematically addressed how the skin immune landscape is altered during such “cryptic” infections.

The skin represents our major barrier to the outside world, reflected by its complex microanatomy. Below an epidermal barrier capable of extensive remodelling in response to traumatic injury and aging lies the dermis, a complex microenvironment comprised of stromal elements and immune cells that lends itself to interrogation by systems level approaches. These have been used successfully to decipher pathways of skin stromal cell development ^11,12^ and how wound healing across the life span is accompanied by an altered inflammatory milieu ^13^. Alteration in skin landscape have similarly been studied in psoriaisis and atopic dermatitis ^14^ and cancer ^15^, promising to contribute to the development of new therapeutic approaches to target these and other skin disorders. Skin-microbiota interactions have received much attention, with implications for normal homeostasis, immune system health and the response to infection ^16,17^. However, how the skin reacts to many common and extrinsic environmental stimuli remains poorly understood.

Ultraviolet radiation (UVR) is important for normal physiology, melanogenesis and vitamin D production ^18^ and a potent environmental modifier of immune function [reviewed in ^18^]. Representing ~9% of total UVR in sunlight ^19^, UVB can suppress primary and anamnestic responses and result in antigen specific tolerance ^20^. UVB exposure has therapeutic value in the treatment of psoriasis via modulation of autoinflammatory pathways^21^, has been shown to affect metabolic programming and senescence in keratinocytes ^22,23^, can drive transcriptional changes associated with photo-aging ^24^ and can induce epigenetic changes during carcinogenesis ^25^). In models of cutaneous leishmaniasis (CL), UVB has varied effects depending on parasite species and/or level of UVB exposure ^26,27^. UVB has also been implicated in the pathogenesis of post kala-azar dermal leishmaniasis (PKDL), an important skin sequela of VL ^5,28–30^. However, the impact of UVB on cryptic skin infections with *Leishmania*, or indeed on any infection, has not been studied previously.

Here, we sought to define the how the immune and stromal cell landscape is remodelled during cryptic skin infection with *L. donovani* and how this remodelling is impacted by UVB exposure. Using single-cell RNA sequencing (scRNA-Seq), spatial transcriptomics and inferential network analysis, we identified key cellular and molecular changes associated with skin infection and demonstrate that effector lymphocytes and macrophages, key players in anti-leishmanial immunity and skin inflammation, occupy discrete cellular niches following UVB exposure. We conclude that UVB leads to a profound re-wiring of immune and stromal cell interaction networks during infection, on par with those reported to be induced by microbiota, and that this may facilitate cryptic infection.

## RESULTS

### Development of a UVB exposure model in C57BL/6J mice

To avoid acute skin damage, we established a low-dose UVB irradiation model in female C57BL/6J mice, wherein flank skin was pre-conditioned with UVB 500J/m^2^ three times a week for three weeks ^31,32^ prior to infection and subsequently throughout early infection. Dermatological, parasitological and immunological endpoints were determined at day 16 post infection (p.i.), by quantitative spectrometry, *in vivo* bioluminescence imaging, histology, flow cytometry and single cell and spatial transcriptomics (**Fig. 1a** and Methods).

**Fig. 1:**
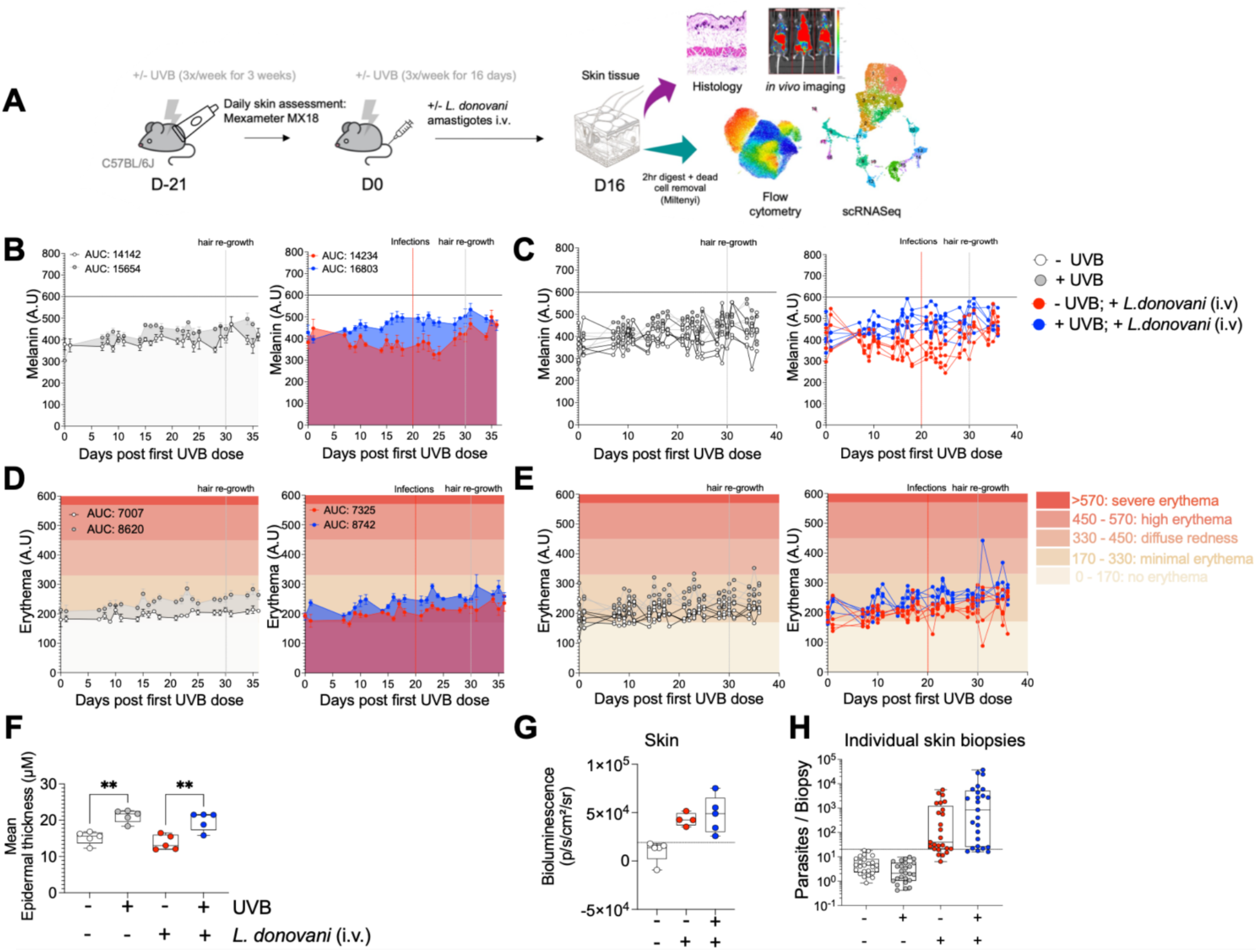
UVB exposure model in control and *L. donovani-* infected C57BL/6J mice. **A)** Schematic diagram detailing UVB exposure, infection and isolation of cells and downstream analysis. **B)** Mean melanin measured in arbitrary units (A.U) for uninfected -UVB and +UVB mice (left panel) and infected -UVB and +UVB mice (right panel). Calculated Area Under the Curve (AUC) is also shown. Horizontal line at 600 on y axis shows melanin values > 600 indicate dark skin; < 600 indicate light skin. **C)** Mean melanin measured in each individual mouse in arbitrary units (A.U). **D)** Mean erythema measured in A.U for uninfected -UVB and +UVB mice (left panel) and infected -UVB and +UVB mice (right panel). AUC is also shown. **E)** Mean erythema measured in each individual mouse in A.U. Symbols as in (D). Erythema scale shown in graduated scale. **F)** Epidermal thickness measured in μM across treatment groups. **G)** Skin bioluminescence imaging across treatment groups. **H)** Parasite load per skin biopsy measured by qPCR. Box plots show the minimum, the maximum, the sample median, and the first and third quartiles. Two-way ANOVA with Tukey’s multiple comparisons test or One-way ANOVA with Tukey’s multiple comparisons test, ** p < 0.01. Data shown are from two independent experiments (n=5 mice per experiment per group).

Mice exposed to UVB (+UVB mice) exhibited minimal changes in melanin (**Fig. 1b and c, and Supplementary Fig. 1a**), minimal erythema ^33^ and only mild epidermal thickening (**Fig. 1f and Supplementary Fig. 1a,b**) compared to non-exposed mice (− UVB mice), irrespective of infection status. Parasites were detectable in the skin and viscera of infected but not uninfected mice, with splenomegaly also evident (**Supplementary Fig. 1c-h**). As previously reported in B6.*Rag2*^−/−^ mice ^5^, parasites in the skin of immunocompetent C57BL/6 showed a patchy distribution and no significant differences were observed due to UVB exposure (**Fig. 1g and h**). Collectively, these data indicate that i) UVB exposure and / or skin parasitism do not lead to clinically significant pathology and ii) that the UVB regimen employed does not directly impact levels of skin parasitism.

### UVB selectively alters skin cell composition during L. donovani infection

We used multi-parameter flow cytometry to generate a high-resolution phenotypic map of immune and stromal cells in uninfected and infected -UVB and +UVB mice (**Fig. 2a and b** and **Supplementary Fig. 2a-c**). In uninfected mice, UVB exposure significantly increased the frequency of CD45^−^Ter119^−^CD31^−^PDPN^+^ stromal cells but decreased the frequency of CD45^−^Ter119^−^CD29^+^ endothelial cells and CD45^−^Ter119^−^ CD4^−^ CD49f^hi^ CD34^+^ hair follicle stem cells (HFSCs). *L. donovani* infection similarly increased the frequency and number of CD45^−^Ter119^−^CD31^−^PDPN^+^ stromal cells compared to uninfected mice, often to a greater extent, but this was dampened by UVB exposure. Furthermore, *L. donovani* infection reduced the frequency and number of endothelial cells and HFSCs, with both effects exacerbated in +UVB mice (**Fig. 2b** and **Supplementary Fig. 2d**). No other significant differences were observed across the remaining stromal cell populations analysed. Amongst immune cells, we noted reductions in frequencies and numbers of CD11b^+^ Ly6C^int^ and CD11b^+^Ly6C^hi^ monocytes in infected +UVB mice compared to infected -UVB mice (**Fig. 2c and d** and **Supplementary Fig. 2d**). No other significant differences were evident across the remaining immune cell populations analysed, although there was a trend towards an increase in T cells and NK cells following infection. Hence, at this level of phenotypic analysis, UVB exposure induces changes in stromal and immune populations that were either mimicked or exaggerated by infection. Superimposing UVB and infection resulted in additional changes, indicative of a complex interplay between these two skin insults.

**Fig. 2:**
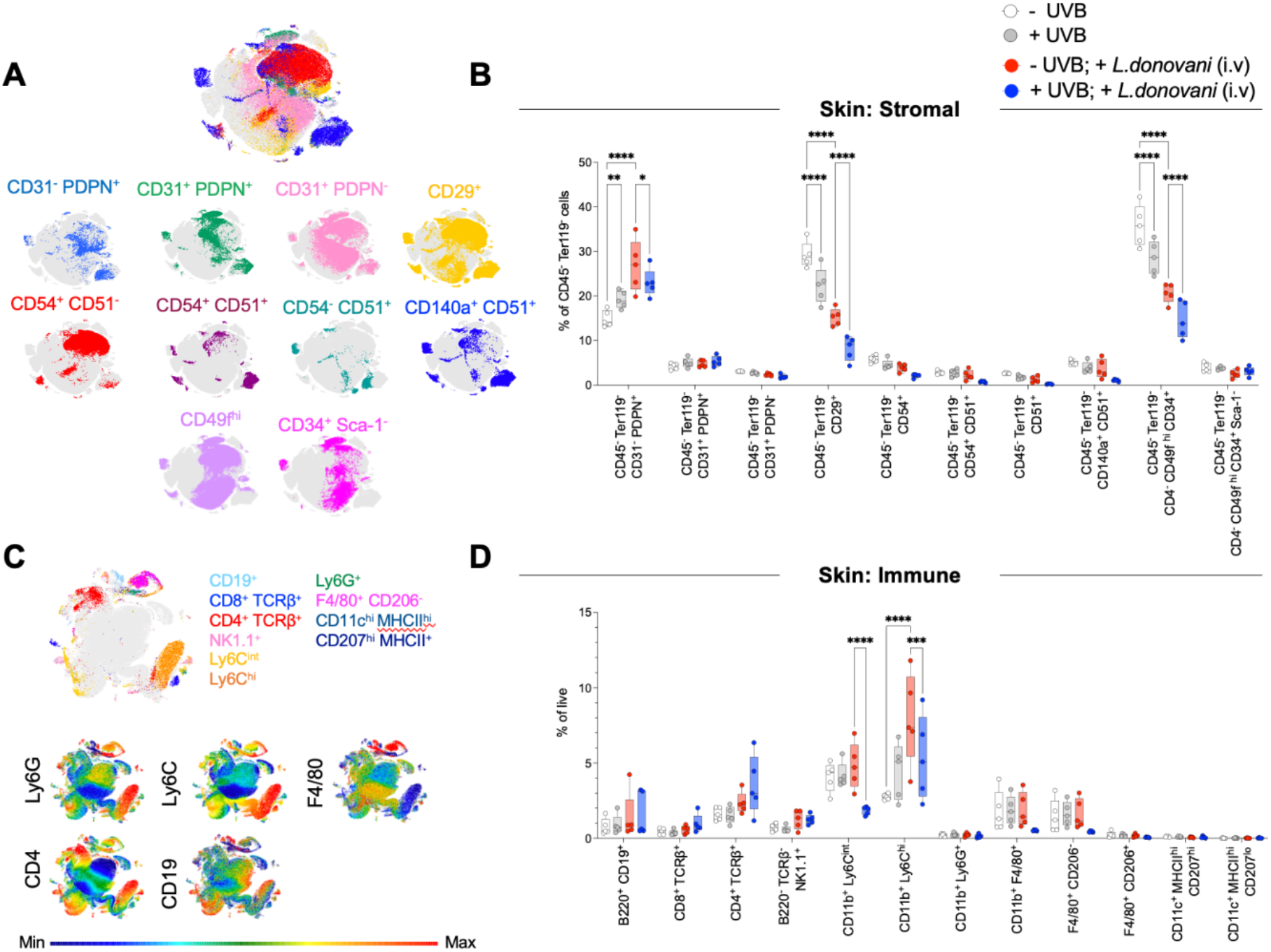
UVB modulates stromal and immune cell populations in the skin. Skin tissue was processed for flow cytometric analysis as described in Methods. **A)** UMAP plots showing individual stromal populations in the skin. **B)** Frequency of stromal cell populations on day 16 p.i. **C)** UMAP plots showing immune cell populations in the skin. **D)** Frequency of immune cell populations on day 16 p.i.. Box plots show the minimum, the maximum, the sample median, and the first and third quartiles. One-way ANOVA with Tukey’s multiple comparisons test; * p < 0.05, ** p < 0.01, *** p < 0.001, ****p < 0.0001. Data shown are representative of two independent experiments (n=4-5 mice per experiment per group).

### scRNA-Seq analysis of the skin cellular landscape during L. donovani infection

To characterise changes in the skin microenvironment more fully, we generated an integrated scRNA-Seq cell atlas from 34,705 cells across all conditions. Using consensus markers ^34,35^, we identified 16 cell clusters based on cluster-specific gene expression (**Fig. 3a**, **Supplementary Fig. 3**, **Supplementary Table 1**). We mapped the spatial distribution of these clusters in healthy non-UVB exposed skin using the 10X Visium platform (**Fig. 3b and c**). Fibroblasts FI and FII represented papillary fibroblasts with a sub-epidermal location. Fibroblasts FIII – FVI were largely restricted to the reticular dermis, muscle and adipose tissues, though FV and FVI also localised to occasional dermal areas populated by macrophages, T cells and NK cells. Macrophages were prominent in the dermis as well as adipose tissue (**Fig. 3c**). We then performed pairwise comparisons across experimental groups to generate ‘response-specific” transcriptional signatures from the scRNA-seq data.

**Fig. 3:**
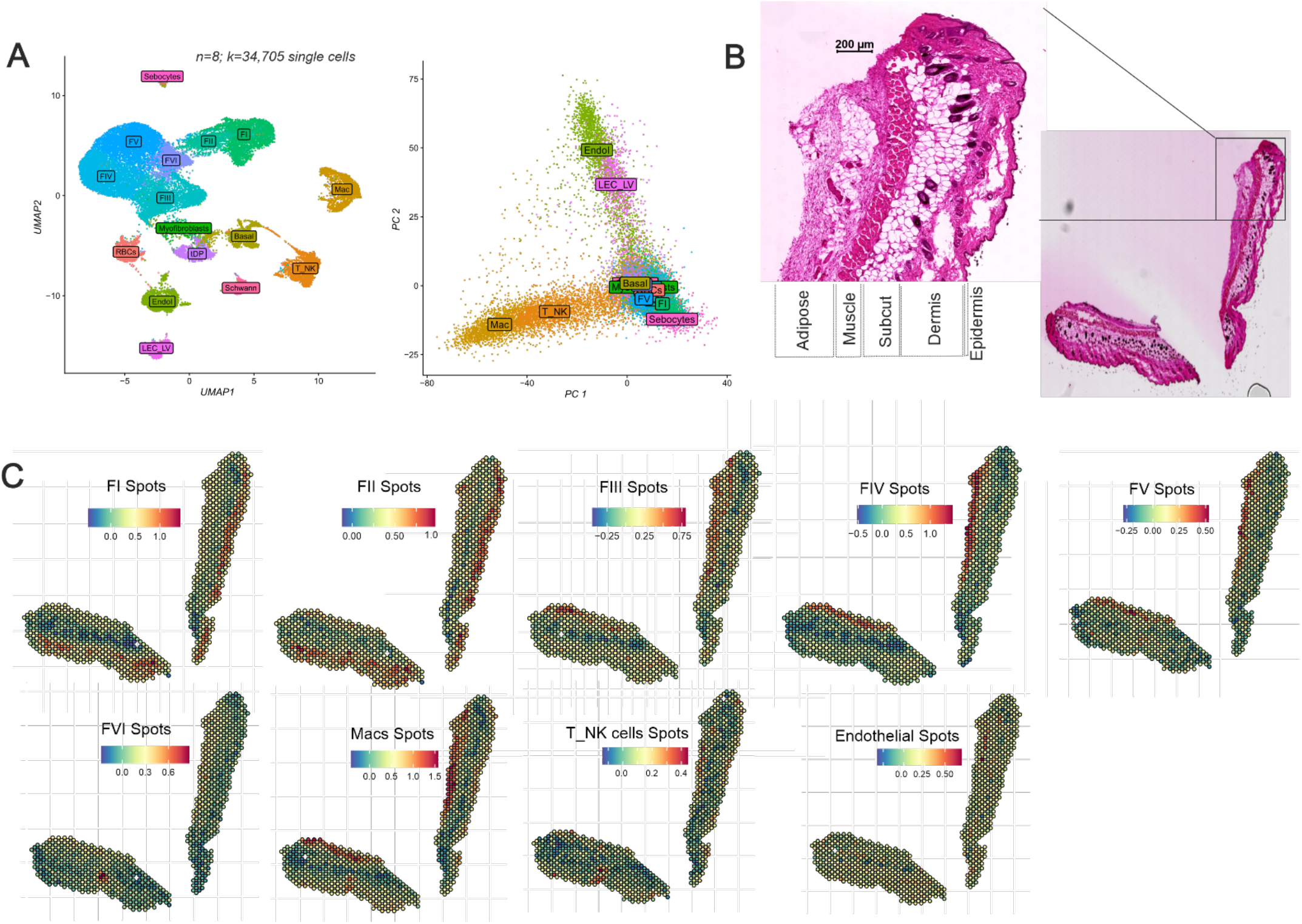
Single cell and spatial transcriptomic diversity of mouse skin. Skin tissue was processed for scRNA-seq as described in Methods. **A)** Scatter plots showing cellular diversity in skin among all groups along with their imputed cell types (left) visualised using UMAP axes. Cell types visualised using the first two principal components (right). **B)** H&E-stained image of skin from -UVB mouse. **C)** Spatial plots showing position of enrichment of cell types (from A) in 55 μm Visium spots on -UVB mouse skin (from B). Data are derived from scRNA-seq analysis of 34,705 cells (k) from 8 mice (n).

#### L. donovani infection induces local transcriptional changes

We first compared *L. donovani* infected and uninfected -UVB mice, providing an unprecedented view of the local response to cryptic infection in the conventional mouse model (**Fig. 4a-c** and **Supplementary Fig. 4a and b**). Infected skin had a greater proportion of FV (*Ptx3, Ccl2, Cxcl1*) and FVI (*Cxcl9, Cxcl1, Gpb2*) fibroblasts and basal cells / keratinocytes (*Sfn, Krt14, Krt17*) compared to uninfected skin (**Fig. 4a**). Conversely, infected skin had a reduced frequency of FI fibroblasts and endothelial cells (*Fabp4*, *Cd36*, *Aqp1*) when compared to uninfected skin. Differential gene expression (DGE) analysis identified major changes to gene expression in each cell population (**Supplementary Fig. 4b**). *Ccl2* was most highly upregulated in T_NK cells, endothelial cells, fibroblast populations FI, FIII, FIV and Schwann cells (**Fig. 4b**), highlighting a potential role for chemokine-mediated cellular recruitment during infection. Importantly *Cxcl1*, involved in inflammasome activation ^36^ was the most significantly upregulated gene in the macrophage cluster (*Retnla, Lyz2, Ccl6, Fcer1g*) and in fibroblasts. *Cxcl2* and *Ccl2* were also both highly upregulated in fibroblast populations FII, FIII, FIV, FV along with lymphatic endothelial cells (LEC_LV; **Fig. 4b and Supplementary Fig. 4b**). Gene set enrichment analysis ^37^ of the top 25 upregulated genes in FV fibroblasts, basal cells, macrophages and T_NK cells indicated an upregulation of IFNγ response genes in FV fibroblasts and macrophages (**Fig. 4c**). Genes involved in TNF signalling via NF-κB were upregulated in FV fibroblasts, macrophages and T_NK cells. Genes upregulated in basal cells were enriched for the p53 pathway (likely associated with stress responses), and pathways associated with coagulation and IL2-STAT5 signalling.

**Fig. 4:**
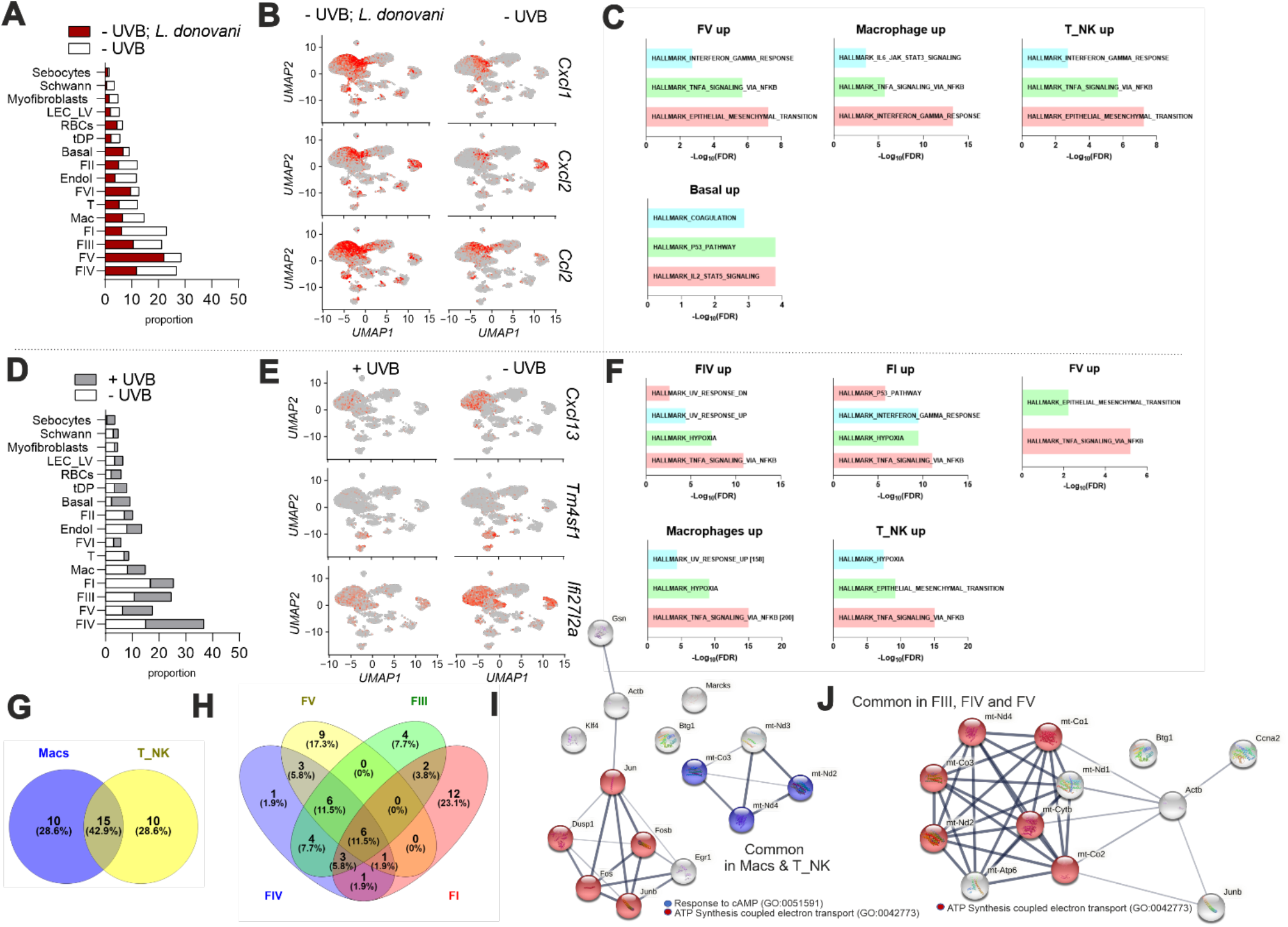
Gene expression changes associated with *L. donovani* infection and UVB exposure. **A)** Cell proportions in uninfected and infected -UVB mice. **B)** UMAP plots for infected and uninfected -UVB mice showing expression of *Cxcl1, Cxcl2* and *Ccl2*. **C)** GSEA enrichment (negative log FDR) for genes upregulated in fibroblast FV, macrophages, T_NK and Basal cells. **D)** Cell proportions in uninfected +UVB vs. -UVB mice. **E)** UMAP plots for uninfected +UVB and -UVB mice showing expression of *Cxcl13, Tm4sf1* and *Ifi27l2a*. **F)** GSEA enrichment for genes upregulated in fibroblasts FIV, FI and FV, macrophages and T_NK cells. **G)** Venn diagram to depict common genes upregulated in macrophages and T cells of +UVB vs. -UVB mice. **H)**. Same as G but for fibroblasts **I)** STRING networks of commonly upregulated genes in macrophages and T_NK cells, coloured by gene ontology terms. **J)** STRING network of commonly upregulated genes in FIII, FIV and FV. Data are derived from scRNA-seq analysis of 23,495 cells for A & B and 17,283 cells for D & E

In addition to *Ccl2* and *Cxcl1*, *Gsn* (gelsolin; an actin-binding protein associated with apoptosis and invasion), *Dcn* (decorin; an extracellular matrix protein secreted by T cells that can inhibit chemokine signalling ^38^) and Mt1 (metallothionein 1; involved in T cell differentiation ^39^) were upregulated in T_NK cells. *Ptx3*, previously identified as a regulator of CD4^+^ T cell responses in CL ^40^ was upregulated in FII and FV fibroblasts as well as other populations (**Supplementary Fig. 4b**). Collectively, these data provide the first overview of the changing immune landscape associated with cryptic *L. donovani* skin infection.

#### UVB exposure reduces CXC-ligand expression and drives metabolic re-programming in immune and stromal cells

Next, we identified changes solely attributable to UVB under these exposure conditions, comparing uninfected +UVB and -UVB mice (**Fig. 4d-j**). Reduced proportions of FI and FII fibroblasts, myofibroblasts and T_NK cells were observed in +UVB mice compared to -UVB mice (**Fig. 4d**), with concomitant alterations in gene expression profile (**Fig. 4e** and **Supplementary Fig. 4c**). Notably, *Cxcl1* (FVI fibroblasts), *Cxcl2* (macrophages, FVI fibroblasts), *Cxcl13* (FIV fibroblast) were downregulated on UVB exposure. *Tm4sf1*, necessary for endothelial migration ^41^ was also down regulated in endothelial cells and basal cells, whereas in macrophages and fibroblasts (FIII, FIV and FV), *Ifi27l2a*, a candidate anti-viral ISG 42, was down-regulated (**Fig. 4e**). Fibroblast populations FII – FVI show increases in *mt-Co3*, a mitochondrial cytochrome c oxidase gene that drives oxidative phosphorylation and has been reported to increase upon UVB exposure ^43^. Furthermore, we noted reduced accumulation of *Rarres2*, *Cyp2f2* and *Angptl1* upon UVB exposure in FI fibroblasts (**Supplementary Fig. 4c**) supporting the hypothesis of metabolic reprogramming of fibroblasts in +UVB compared to -UVB mice, and consistent with a recent study of human fibroblasts exposed to UVB ^44^.

GSEA analysis of upregulated genes in +UVB mice highlighted TNF signalling and hypoxia in FIV and FI fibroblasts (**Fig. 4f**) and Macrophages and T_NK. Genes associated with UV response were enriched in FIV and FV fibroblasts and macrophages. To understand if UVB exposure induced a set of core genes in immune cells and fibroblasts, we looked at the intersection of the top 25 upregulated genes in macrophages and T_NK cells (**Fig. 4g**) and in fibroblasts (**Fig. 4h**). We found that 43% of upregulated genes were common to both macrophages and T_NK cells and related to oxidative phosphorylation and cAMP (cyclic AMP) stimulus (**Fig. 4i**). Genes exclusively upregulated in macrophages were enriched for NADH dehydrogenase activity and protein folding. Conversely, uniquely regulated T_NK cell genes were enriched for extracellular matrix constituents (**Supplementary Fig. 4d**). Among fibroblast populations, genes upregulated upon UVB exposure also overlapped (**Fig. 4h**), with common genes enriched for electron transport chain (**Fig. 4j**). FI fibroblasts had ten distinct genes enriched for MHC class I presentation (**Supplementary Fig. 4d**).

#### UVB exposure modifies the skin response to L. donovani infection

Having established the baseline effects of UVB exposure, we next sought to examine how infection-associated gene expression patterns differed following UVB exposure (**Fig. 5**). Consistent with flow cytometric analysis (**Fig. 2**), we found an increase in the proportion of T_NK cells in the skin of infected +UVB mice compared to uninfected +UVB mice (**Fig. 5a**). *Ccl5, Gzma and Ifng* were amongst the top upregulated genes in T_NK cells in infected +UVB mice (**Fig. 5b** and **Supplementary Fig. 5a**). To further define these cells, we sub-clustered the T_NK cluster to reveal 7 sub-clusters, defined as NK cells (*Nkg7*, *Gzma*), naïve T cells (*Il7r*), Th1-like (*Ifng, Ccl5*) and Th2-like (*Il5*, *Gata3*) TCRαβ T cells, TCRγδ (gDT) T cells and proliferating T cells (*Mki67*, *Tubb5*), along with a small contaminating subset of mast cells (*Mcpt4*, *Cma1*) (**Fig. 5c-e** and **Supplementary Table 2**). *Ifng* transcripts were largely restricted to Th1-like CD4^+^ cells and this sub-cluster was over-represented amongst T_NK cells in skin of infected +UVB mice compared to uninfected +UVB mice, largely at the expense of naive T cells, TCRγδ T cells, and proliferating T cells (**Fig. 5e** and **Supplementary Table 2**).

**Fig. 5:**
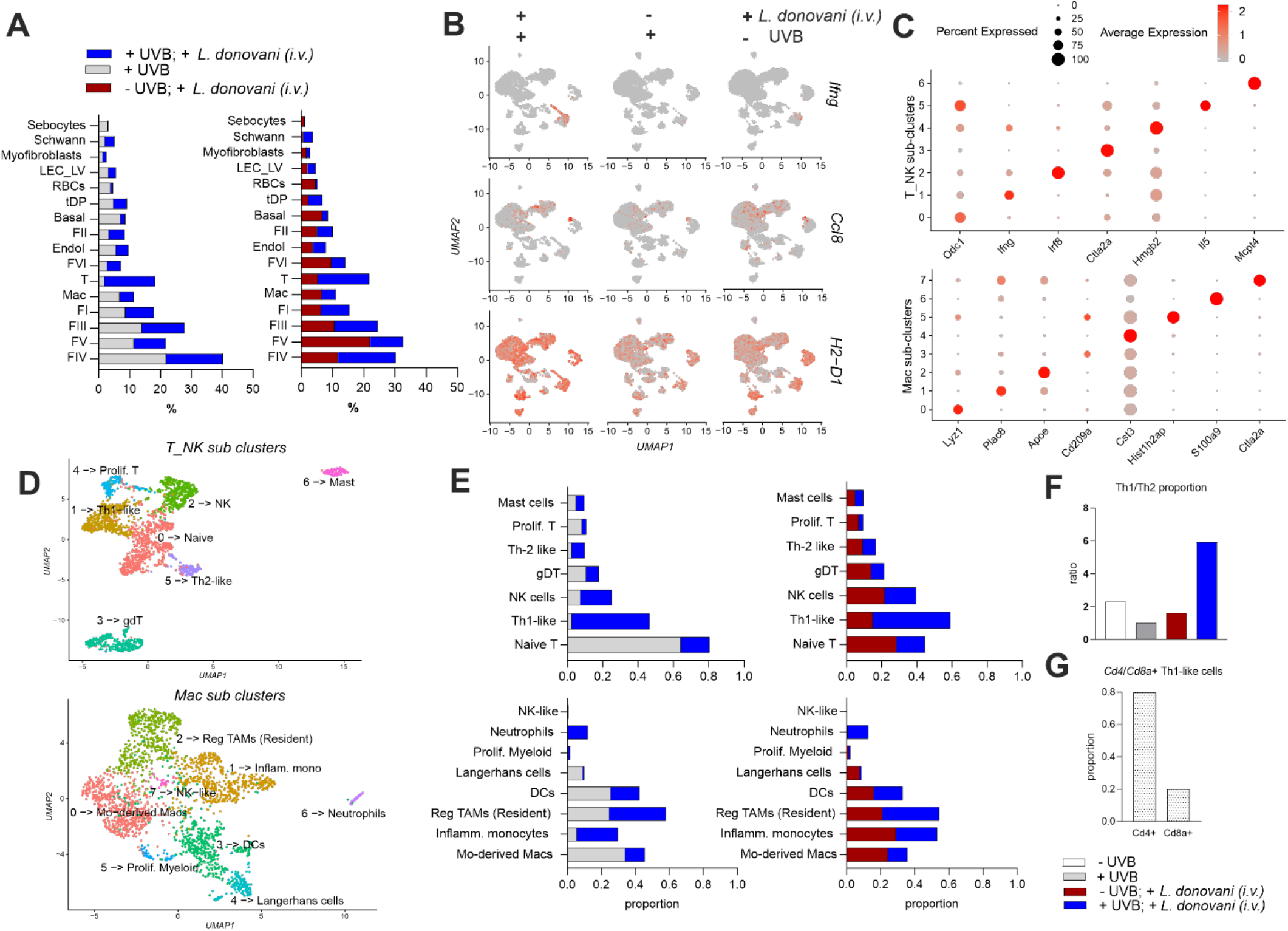
UVB pre-exposure affects immune populations differentially upon *L. donovani* infection. **A)** Cell proportions shown for infected and uninfected +UVB mice (left) and infected +UVB vs. -UVB mice (right). **B)** UMAP plots to show gene expression of *Ifng, Ccl2, Ccl8, H2-D1* across comparisons shown in (A). **C)** UMAPs showing sub-clusters within original T_NK (above) and macrophage (below) clusters. **D)** Dot plots showing the top genes expressed in T_NK (above) and macrophage (below) sub-clusters. Cluster numbers refer to populations shown in (C). **E)** Cell proportions of imputed sub-clusters in (C) and (D), comparing infected and uninfected +UVB mice (left) and infected +UVB vs. -UVB mice (right). **F)** Bar plot shows ratio of Th1-like and Th2-like cells across all groups. **G)** Bar plot showing distribution of *Cd4*^+^ and *Cd8*^+^ cells within the Th1-like population. Data are derived from scRNA-seq analysis of 24,109 cells for A, B whereas sub-cluster analysis for T_NK and Mac are based on 2,274 and 2,362 cells respectively.

Likewise, macrophages were sub-clustered to reveal Langerhans cells (*Cd207*, *Cts3*), neutrophils (*S100a9, S100a8*), monocyte-derived macrophages (*Cd14*), dendritic cells (*Cd209*), inflammatory monocytes (*Cd14*, *Lyz2*, *Plac8*, *Ly6c*), regulatory/resident macrophages (*Adgre1*, *Apoe*, *Mrc1, C1q, Retnla*), proliferating myeloid cells (*Mki67*) and a minor population of NK-like cells (**Fig. 5c and d** and **Supplementary Table 2**). The proportions of each population varied across treatment group, with infected +UVB mice showing substantially raised neutrophils and monocytes compared to uninfected +UVB mice largely at the expense of monocyte-derived macrophages, DCs and Langerhans cells (**Fig. 5e** and **Supplementary Table 2**). A 2 log-fold increase in *Ccl8*^+^ was observed in macrophages in infected +UVB mice compared to uninfected +UVB mice (**Fig. 5b** and **Supplementary Fig. 5a**) associated largely with *Apoe*^+^*Mrc1*^+^ *C1qb^+^* cells, a phenotype reminiscent of murine lipid-associated tumour-associated macrophages (LA-TAMs) / immune regulatory (Reg-) TAMs ^45^. Based on higher abundance of *Adgre1* mRNA, these cells may also be resident macrophages of fetal origin ^46^.

Multiple cell populations showed an increase in MHCI genes, this being most notable in fibroblasts in infected +UVB mice (**Fig. 5b** and **Supplementary Fig 5c**). Whereas mitochondrial cytochrome genes related to oxidative phosphorylation were upregulated in fibroblasts in uninfected +UVB mice compared to uninfected -UVB mice (**Fig. 4h and j**), many of these genes were downregulated when comparing infected +UVB mice to uninfected +UVB mice (**Supplementary Fig. 5**). This suggests that while UVB exposure increases steady state expression in fibroblasts, subsequent infection may downregulate such responses. For example, downregulation of *mt-Co3* is seen across most cell types (**Supplementary Fig. 5**). The long non-coding RNA *Malat1*, that participates in UVB-induced photo-aging in fibroblasts ^47^, was in the top 5 upregulated transcripts in multiple cell types when comparing infected +UVB mice and uninfected +UVB mice (**Supplementary Fig. 5**). Hence, concurrent cryptic infection can modulate effects associated with UVB exposure.

We next directly compared transcriptional responses in infected +UVB and -UVB mice to delineate the differences more clearly between immune status in these two models (**Fig. 5a, b and e** and **Supplementary Fig. 5c**). This analysis confirmed expansion of T cells and FIV fibroblasts and a reduction of FV fibroblasts in infected +UVB vs infected -UVB mice. GSEA of the top 25 upregulated genes highlighted predominantly pro-inflammatory pathways (**Supplementary Fig. 5e**) accompanied by enhanced *Ifng* expression by Th-1 like cells (**Fig. 5b**). Indeed, the ratio of Th1-like to Th2-like cells was ~5 fold higher in infected +UVB mice compared to infected -UVB mice (**Fig. 5f**). Within Th-1 like cells, the ratio of *Cd8a^+^* to *Cd4^+^* cells was 1:4 (**Fig. 5g**). Strikingly, the *Ccl8*^+^ macrophage signature was retained in this comparison, confirming the increase in regulatory / resident macrophages (**Fig. 5b)**. *Apod* was also increased in fibroblasts FIII and FVI (**Supplementary Fig. 5a and d**). Collectively, these direct comparisons indicate that under conditions of UVB exposure, the skin response to cryptic *L. donovani* infection is skewed towards the expression of pro-inflammatory gene pathways and effector T and NK cell potential.

*Cxcl9* and *Cxcl12*, associated with regulating inflammation, leucocyte recruitment and tertiary lymphoid structures (TLS) formation in chronic inflammation and cancer 48, were upregulated in multiple cell populations (notably endothelial cells and FIII fibroblasts) in infected +UVB compared to uninfected +UVB mice (**Supplementary Fig. 5b**). Given their role in the formation of TLS ^48^ and skin inflammation ^49^ and our flow cytometry data indicating changes in the abundance of CD31^−^PDPN^+^ stromal cells (**Fig. 2**), we further examined the heterogeneity of *Pdpn* expression across all groups. As anticipated, *Pdpn* transcripts were associated to a greater or lesser extent with all fibroblast populations, with somewhat heightened expression in FII, FV and FVI fibroblasts after infection. UVB exposure had a deleterious effect on *Pdpn* expression on FV fibroblasts and LEC_LV’s (**Supplementary Fig. 6a**).

**Fig. 6:**
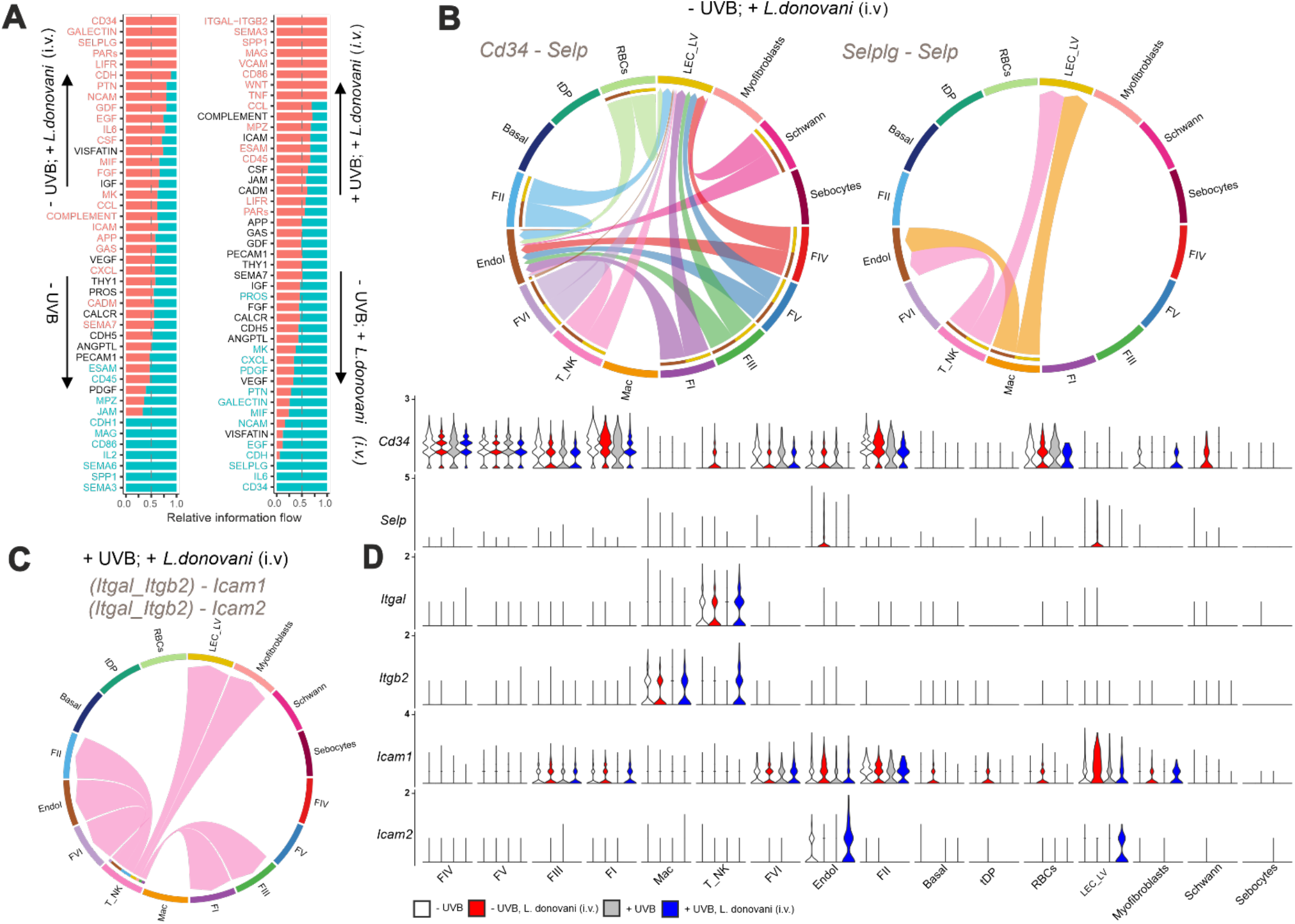
UVB exposure re-wires immune and stromal cell cross talk. scRNA-seq data was analysed using CellChat as described in Methods. **A)** Pairwise comparison of information flow between inferred signalling networks, with pathways in red more enriched in groups indicated with upward arrow and vice versa. **B)** Chord plots show directional flow of information (arrow heads indicate receiving cells/receptors) between cell types for “*Cd34–Selp*” (left) and “*Selplg-Selp*” (right) interactions in infected -UVB mice. **C)** Same as B, but for *(Itgal+Itgb2)-Icam1/2* interactions in infected +UVB mice. **D)** Violin plots showing normalised gene expression for each of the cell types split across comparison groups. Data are derived from scRNA-seq analysis of 34,705 cells for A and D whereas B and C are derived from 12,899 and 4,523 cells respectively.

#### Modification of immune and stromal cell circuits following UVB exposure

We used CellChat ^50^ to identify “secreted signalling” and “cell-to-cell contact” pathways that provide the framework for cross-talk between immune and stromal cells (**Fig. 6, Supplementary Fig. 6b and c**). Amongst the latter, we identified a prominent role for CD34 / p-selectin (*Cd34*-*Selp*) and P-selectin glycoprotein 1 (SELPLG) / p-selectin (*Selplg-Selp*) pathways in the skin of infected -UVB mice. However, these interactions were absent in infected +UVB mice (**Fig. 6a**). We then identified the top ligand-receptor pairs that contribute to signalling in infected +UVB and -UVB mice. *Cd34-Selp* interactions scored highest among the top 5 exclusive interactions in infected -UVB mice (**Supplementary Fig. 6b**). Except for basal cells, tDPs (telogen dermal papilla), myofibroblasts and sebocytes, all cells (including macrophages and T_NK cells) signal to endothelial cells and LEC_LVs via the *Cd34-Selp* axis (**Fig. 6b**). We confirmed *Selp* expression in endothelial cells and LEC_LVs occurred only in infected -UVB mice (**Fig. 6d**). CellChat probabilities also indicated that *Selplg-Selp* interactions occur between macrophages and T_NK cells and endothelial cells / LEC_LV again only in infected -UVB mice (**Fig. 6a and b**). This likely reflects the loss of *Selp* mRNA in endothelial and LEC_LV after UVB exposure (**Fig. 6d**). In contrast to the role of p-selectin in infected -UVB mice, the prominent cell-to-cell contact interactions in infected +UVB mice involved T_NK cell signalling via *Itgal/Itgb2 (*LFA1*) - Icam1* and *Itga /Itgb2* - *Icam2* to myofibroblasts, FII, FVI, FI, FIII fibroblasts, endothelial cells and LEC_LVs (**Fig. 6a and c**).

To further explore these interactions, we identified 6 sub-clusters of endothelial cells (**Fig. 7a and Supplementary Fig. 7a and b)**, with *Selp* largely restricted to endothelial subcluster 1, bearing a gene signature indicative of cytokine activation (E-selectin, GM-CSF, von Willibrand Factor, aquaporin-1) (**Fig. 7b**). *Selp* mRNA was also abundant in a subpopulation of LEC_LVs (**Supplementary Fig. 7c-e**).

**Fig. 7:**
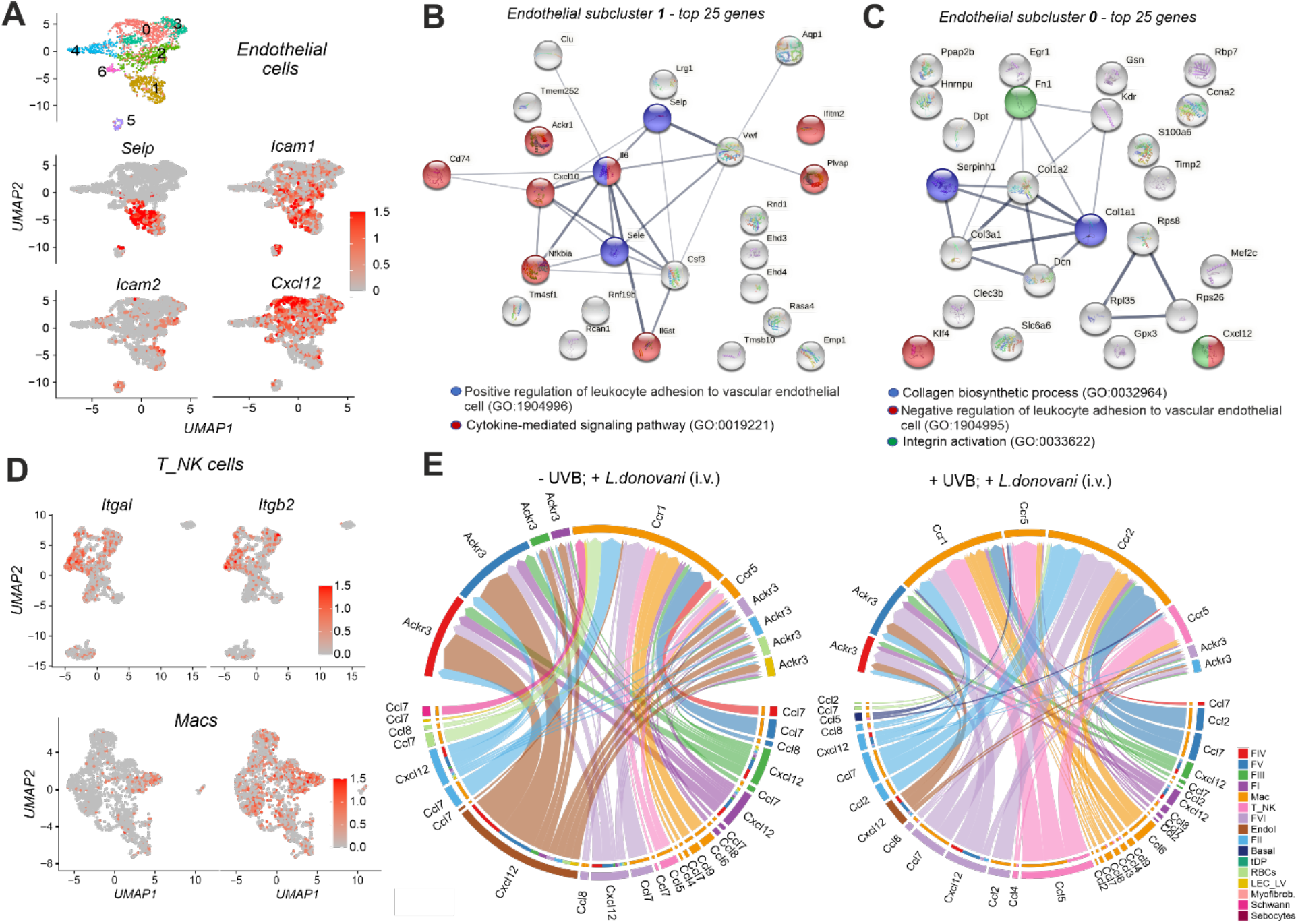
Expression and sub-cluster localisation for immune and stromal cell signalling molecules. **A)** Cluster memberships (colours) on sub-clustered endothelial cells along with *Selp, Icam1, Icam2* and *Cxcl12* expression shown on UMAP. **B)** STRING networks of commonly upregulated genes in endothelial subcluster 1 coloured by gene ontology terms. **C)** Same as B but for endothelial subcluster 0. **D)***Itgal* and *Itgb2* expression shown on UMAP plot of T_NK cells (above) and Macrophages (below). **E)** Chord diagram representing *Ccl-* and *Cxcl-* mediated networks in infected -UVB vs. infected +UVB mice. Colours indicate individual cell type and are indicated in the legend. Data for A) and D) are derived from scRNA-seq analysis of 1894 endothelial, 2274 T_NK, 2362 Mac cells whereas for E data is derived from 12,899 and 4523 cells for infected -UVB and infected +UVB respectively.

Interestingly, *Icam2* expression in the endothelial subclusters showed no cluster-specificity whereas *Icam1* was found to be expressed in subclusters 1 and 2 (**Fig. 7a**). In LEC_LVs, *Icam2* was broadly expressed, whereas *Icam1* was restricted to subcluster 0 (**Supplementary Fig. 7e**). *Itgal* and *Itgb2* transcripts that contribute to increased signalling in infected +UVB mice were identified in the Th1-like and NK sub-clusters of T_NK population (**Fig.s 5d and 7d**). *Itgb2* expression was broadly expressed in macrophages whereas *Itgal* was upregulated in macrophage subcluster 1/inflammatory monocytes (**Fig.s 5d and 7d**).

We next explored secreted signalling pathways (**Fig. 7e**) given our earlier observation that infection induces a diverse chemokine response (**Fig. 4b, and Supplementary Fig. 6**) and the results from ligand-receptor predictions (**Fig. 6a**). While *Cxcl12*-*Ackr3* interactions were prominent in both infected +UVB and -UVB mice (**Supplementary Fig. 6b**), endothelial-led CXCL12-ACKR3 interactions were predicted to be markedly reduced in infected +UVB mice. Notably, *Cxcl12* was the top marker gene for endothelial subcluster 0 (**Fig. 7a, 7c)** and the proportion of endothelial cells in this sub-cluster was reduced two-fold in infected +UVB mice compared to infected -UVB mice (**Supplementary Fig. 7b**). Of note, CCL2-CCR2 interactions were not predicted in -UVB mice (**Fig. 7e**, left), despite the prominence of a *Ccl2* signature in the skin (**Fig. 4b**), suggesting receptor availability is limiting. In contrast, this ligand-receptor pathway was observed in infected +UVB mice, associated with the greater expression of *Ccr2* by cells in the macrophage cluster (**Fig. 7e**, right). Finally, given the prominent recruitment of *Ifng^+^Ccl5^+^* Th1-like cells in infected +UVB mice and the finding that CC chemokine pathways also appeared differently affected by UVB exposure, we examined CXC- and CC-chemokine circuits. We found that T cells in infected +UVB mice communicated with macrophages through *Ccr5*, in addition to the usage of *Ccr1* seen in infected -UVB mice (**Fig. 7e**). Furthermore, the main pathways of chemokine interaction between stromal cells themselves and stromal cells and immune cells were significantly different in infected +UVB compared to -UVB mice (**Supplementary Fig. 6b**). Collectively, these data indicate that UVB precipitates a wholescale re-shaping of the inter-cellular networks established during cryptic *L. donovani* skin infection.

Finally, to understand the spatial relationships between cell types identified in our scRNA-seq data, we used Cell2location ^51^ to assign cell type abundances to individual Visium spots (**Supplementary Fig. 8** and Methods). In infected -UVB mice, examination of the correlation between cell abundances indicated the presence of specific cellular niches (**Fig. 8a**). We used these correlations as distances to build graphs and calculated minimum spanning tree-based clusters. In infected -UVB mice, Th1 cells were predicted to most closely associate with regulatory / resident macrophages and monocyte-derived macrophages and FIV fibroblasts (**Fig. 8b**), an associated also apparent when the cell type signatures were mapped back to the Visium data (**Fig. 8c and Supplementary Fig. 8b and 9**). FIII fibroblasts were most closely associated spatially with Endo1 endothelial cells, likely interacting via the CellChat-predicted *Cd34*-*Selp* pathway (**Fig. 8b and Fig. 6b**). Applying the same approach to infected +UVB mice, we found that Th1 cells were now predicted to be more closely associated with Endo1 and Endo2 endothelial cells and spatially separated from both regulatory/ resident macrophages and monocyte-derived macrophages (**Fig. 8d-f and Supplementary Fig. 8b and 9**). As only Th1 cells (in the T_NK population) express *Itgb2* (**Fig.s 5d, 6d** & **7d**) and both Endo1 and Endo2 cells are *Icam1*^hi^ (**Fig. 7a**), this analysis further corroborates our earlier CellChat predictions (**Fig. 6c**). Thus, UVB exposure remodels the cellular niches that contain key players associated with anti-leishmanial immunity and the regulation of inflammation.

**Fig. 8:**
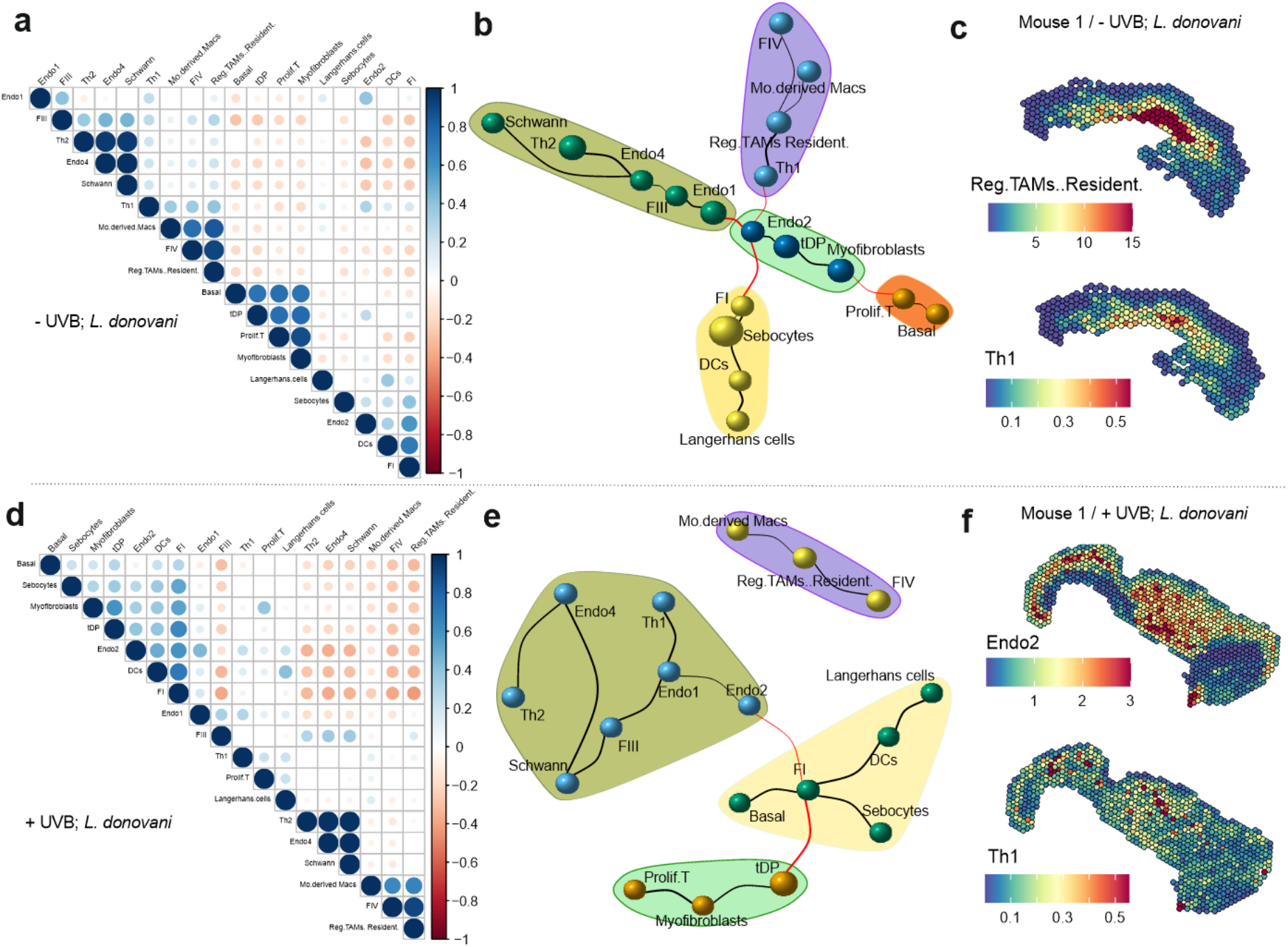
UVB and infection can modify cellular landscapes in skin. **A)** Correlation plots of cell abundances calculated per spot using Cell2location for infected -UVB mice. **B)** Clusters imputed after calculating minimum spanning tree of graphs derived from connecting cells based on their pair-wise distance (1-correlation). Black and red lines indicate connections within and between clusters and the thickness indicates distance. **C)** Spatial plot showing the location of regulatory TAM / resident macrophages with Th1 cells for representative *L. donovani* infected -UVB mouse. **D-E)** Same as A) and B) but for *L. donovani* infected +UVB mouse. **F)** Same as C) but for Endo2 and Th1 cells in representative *L. donovani* infected +UVB mouse. Calculations based on 2101 spots for -UVB and 2985 spots for +UVB mice with each spot measuring 55 micronfrom (n=3 mice per group). Spatial images for additional mice are shown in Supplementary Fig. 8 and 9.

## DISCUSSION

The molecular and cellular characterisation of immune and stromal components of the mouse skin during homeostasis and repair has been reported in great detail ^34,35,50,52–57^, but how these are modified during cryptic infection by trypanosomatid parasites and how such responses might be affected by UVB were previously unknown. Here, we generated a transcriptional atlas of skin immune and stromal populations at rest and after *L. donovani* infection, in the presence and absence of UVB exposure. UVB was found to profoundly alter networks of intercellular communication, associated with heightened T cell activation and proinflammatory cytokine production. Our data provide new insights into i) how the skin microenvironment is altered during cryptic infection with an important human pathogen and ii) extend our understanding of the impact of UVB as an environmental modifier of immunity.

In conventionally housed *L. donovani* infected C57BL/6J mice, skin infection follows a previously described “patchy” distribution (Doehl et al., 2017) without overt clinical pathology, leading to cryptic infection at this site. Nevertheless, we observed significant changes in skin stromal and immune compartments indicative of sub-clinical inflammatory processes when comparing infected vs uninfected mice. Cellular communication in infected mice was dominated by fibroblast – endothelial cell cross talk, mediated through p-selectin signalling and there was an influx of CD11b^+^Lyc6^int^ and CD11b^+^Ly6C^hi^ inflammatory monocytes. Given the concurrent changes in the bone marrow myelopoiesis driven by visceral *L. donovani* infection ^58^, it is likely that these monocytes are direct emigrants from the bone marrow ^59^. PDPN^+^ fibroblastic stromal cells also increased in number and frequency yet despite increased *Cxcl1*, *Cxcl2* and *Ccl2* expression (reminiscent of studies in wounded skin; 35, we did not observe a change in skin T cell abundance. Skin T cells detected in infected mice also had limited effector function suggesting that in mice with concurrent systemic VL, the skin inflammatory response remains somewhat muted.

Stromal cell - endothelial cell interactions are key to the regulation of inflammation ^60^ and a common perivascular fibroblast sub-population expressing SPARC, COL3A1 and POSTN and with pro-inflammatory potential has been identified across multiple human inflammatory diseases ^61^. Such cells bear similarities to the FVI fibroblasts described here and our data represent the first to describe pathways regulating fibroblast-endothelial cell communication in infected skin and to identify the extent of CD34-p-selectin interactions. Selectins have previously been shown to play roles in wound healing ^62–65^ and CD34 is a highly glycosylated transmembrane pan-selectin protein ligand expressed by multiple sub populations of fibroblasts in steady state skin ^34^, in the tumour microenvironment ^66^ and during infection (this manuscript). In contrast to CD34, PSGL-1 is the main leucocyte-expressed ligand of p- and e-selectin ^67^ and we show here that this represents the main ligand in infected -UVB mice for p-selectin-mediated cross talk between T cells and macrophages and endothelial cells / LEC_LV.

Two features stand out from our comparative analysis of infected -UVB and +UVB mice. First, in contrast to the low effector capacity of T cells in infected -UVB mice, Th1-like effector capacity mediated by both CD4^+^ T cells and NK cells was augmented in infected +UVB mice. Paralleling observations made in autoimmune inflammatory diseases ^61^, we interpret these data to suggest that UVB, either alone or in combination with other changes associated with infection, leads to the development of pro-inflammatory stromal cells that promote effector function in skin-infiltrating T cells during *L. donovani* infection. Second, in contrast to infected -UVB mice, where selectins and CXCL12 play prominent roles, inter-cellular communication in infected +UVB mice shows dramatic shifts in chemokine-chemokine receptor bias and towards the use of integrin signalling pathways. These changes appear linked both to a UVB-associated down-regulation of *Selp* expression, loss of *Cxcl12*-expressing endothelial cells and an increased influx of effector CD4^+^ Th1 T cells and NK cells. Spatial mapping of these cell populations demonstrates the existence of discrete cellular niches in the skin of infected -UVB and +UVB mice, significantly extending our previous analysis describing the “patchiness” of the skin immune landscape during infection ^5,68^. Strikingly, under conditions of UVB exposure, we found that effector Th1 cells were located in separate cellular niches to those containing regulatory / resident macrophages and monocyte-derived macrophages. We have previously characterised skin parasite distribution in B6.*Rag2*^−/−^ mice, identifying that ~50% of parasites reside within cells expressing CD206 (ManR) a marker of alternate activation and regulatory phenotype 68. Studies using *L. major* (Seidman) have also defined parasitism of resident / regulatory macrophages as a key event in the establishment and maintenance of persistent infection ^69^. In the current study, we were able to detect skin *L. donovani* parasites by PCR and bioluminescent imaging in immunocompetent C57BL/6 mice, but the limited numbers have made detailed analysis of host cell preferences challenging. Given the paucity of parasites detectable in skin and their patchy distribution, it is likely that some changes we have observed are influenced at least in part by the systemic immune response to infection (rather than the local response in isolation or by parasites *per se*). An analogous situation is observed in patients with systemic sclerosis, where skin *SELP* and *CCL2* transcript abundance strongly correlates with the severity of interstitial lung disease ^70^. Nevertheless, we hypothesize that the spatial segregation between Th1 cells and regulatory / resident macrophages may explain why host resistance is not improved by UVB exposure, despite heightened effector T cell recruitment, a situation that would both favour parasite persistence and set the scene for immunopathology.

PKDL is an important skin complication that often follows treatment for VL ^71^ but understanding of PKDL pathogenesis and the development of new treatments has been hindered by the lack of a pre-clinical model. We and others have speculated based on the clinical pattern of disease development that UVB may play a role in PKDL pathogenesis ^5,28–30^. Examining the role of UVB in this study was in part driven by a desire to develop a model of PKDL but the current study was not designed primarily for that purpose. Whilst UVB enhances the pro-inflammatory environment thought to also underpin PKDL pathogenesis ^72^, clinical symptoms of PKDL did not occur in mice under these conditions of UVB exposure. Two future modifications are likely to be required. First, in humans PKDL usually emerges after recovery from the systemic immunosuppressive state generated by VL ^72^. Although systemic immunosuppression during experimental VL is less pronounced than in human disease, it may nevertheless serve to limit skin inflammation. Inducing systemic parasite clearance through drug treatment would therefore appear an appropriate next step. Second, mouse and human keratinocytes may respond differently to UVB 73-75, requiring the use of genetically-manipulated mouse models to achieve more precise disease positioning.

Our study has some additional limitations. The UVB regimen adopted here balances a requirement for pre-conditioning with the avoidance of acute skin damage, the confounding effects of hair regrowth and the practicalities of animal husbandry and so is not fully reflective of natural UVB exposure in disease endemic countries. We used only female mice to avoid confounding skin inflammation that often results from male aggression and additional studies will be required to determine whether the differences we observe here are sex-dependent.

In conclusion, we have clearly demonstrated the impact of UVB as an environmental modifier of local immune responses during cryptic skin infection by *L. donovani*, altering networks of inter-cellular communication and generating spatially disparate niches within the skin immune-stromal cell landscape. In addition to providing an unparalleled view of the skin response to infection, our data more broadly highlight the importance of considering UVB exposure during the development of translational models for drug and vaccine development against cutaneous infections.

## ACKNOWLEDGEMENTS

The authors would like to acknowledge their gratitude to Karen Hogg, Graeme Park, Sally James, Lesley Gilbert and Katherine Newling (University of York Biosciences Technology Facility) for support with flow cytometry and scRNA-seq analysis, staff in the University of York Biological Services Facility for animal husbandry and Bruce Branchini and colleagues (Department of Chemistry, Connecticut College) for the kind gift of the “red-shifted” luciferase gene. Generation of luciferase-expressing *L. donovani* was funded by the Wellcome Trust (104976/Z/14/Z to E.M. and Jeremy C. Mottram). Graphical abstract created with BioRender.com. This work was funded by Wellcome Trust Investigator Awards to PMK (WT104726, WT224290).

## AUTHOR CONTRIBUTIONS

Conceptualisation, M.M.O, MC and P.M.K; Methodology, M.M.O and P.M.K; Formal Analysis, M.M.O and S.D; Investigation, M.M.O, S.D, K.V.B, H.A, N.B, E.M, N.S.D, G.F.R, E.M, S.J and L.G; Resources, P.M.K, H.A, N.B and D.P.M; Writing – Original Draft, M.M.O and S.D; Writing – Review and Editing, M.M.O, S.D, K.V.B, E.M, M.O and P.M.K; Visualisation, M.M.O and S.D; Supervision, P.M.K; Project Administration, M.M.O and P.M.K; Funding Acquisition, P.M.K

## INCLUSION AND DIVERSITY

We support inclusive, diverse and equitable conduct of research.

## MATERIALS AND METHODS

### KEY RESOURCES TABLE

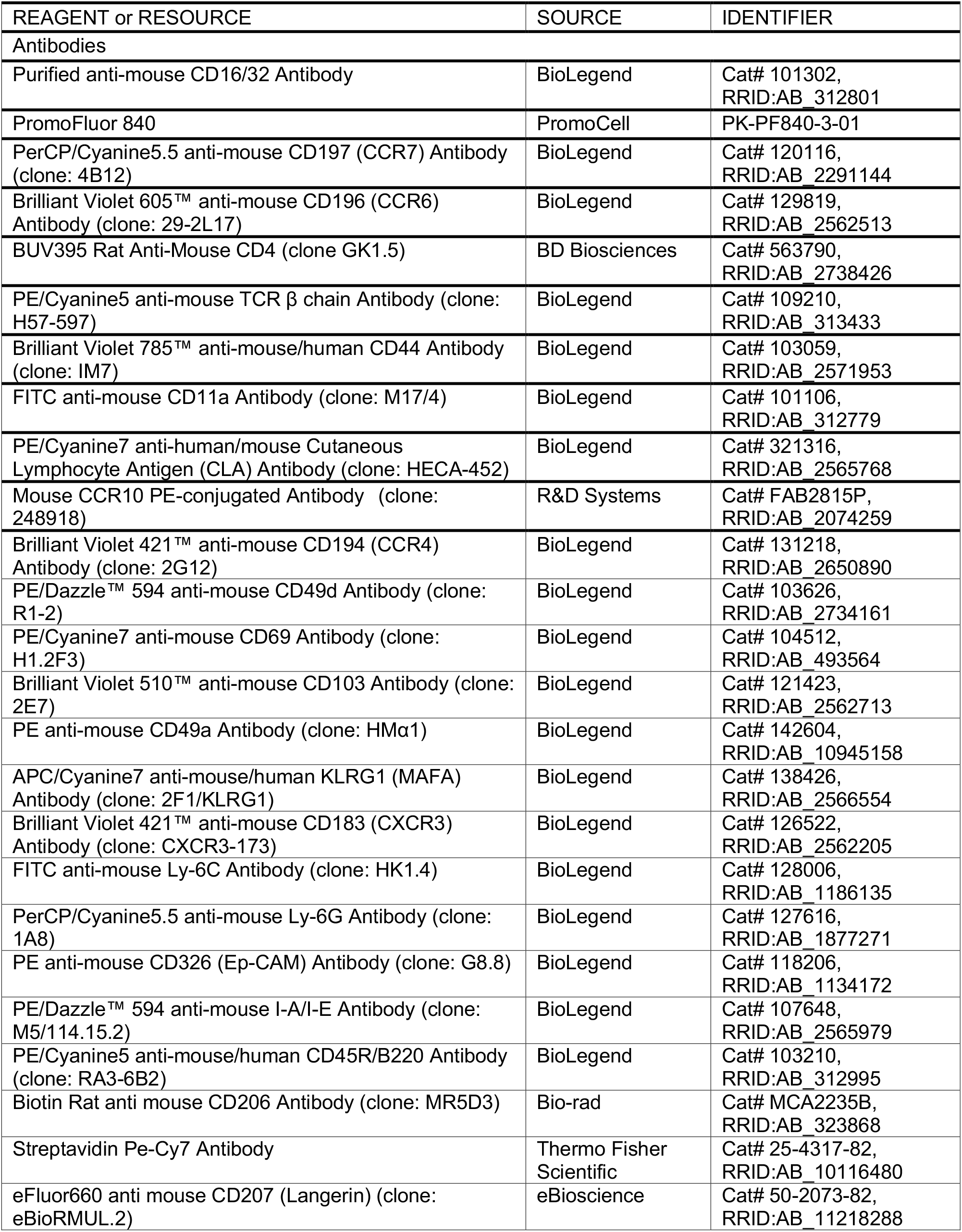
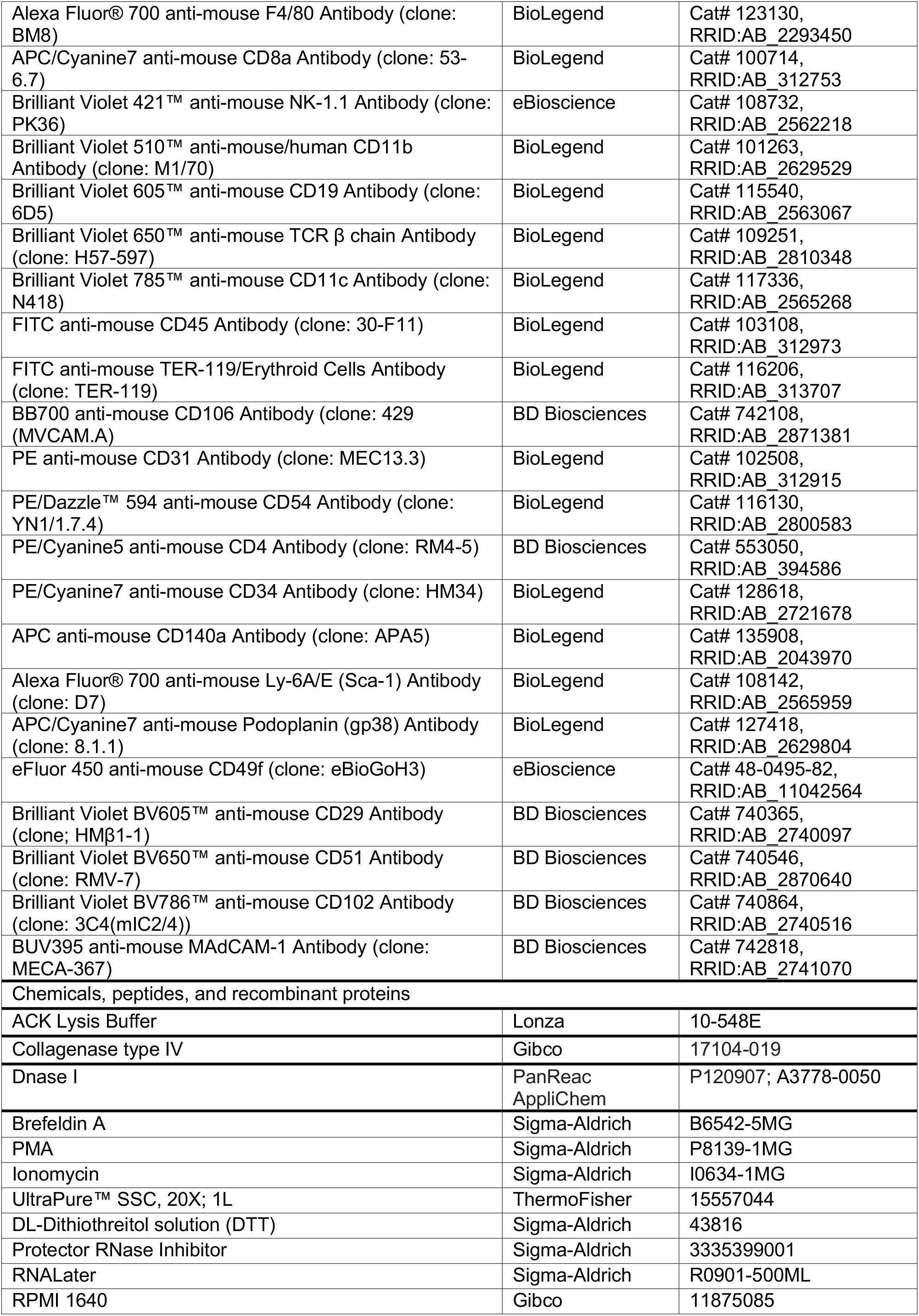
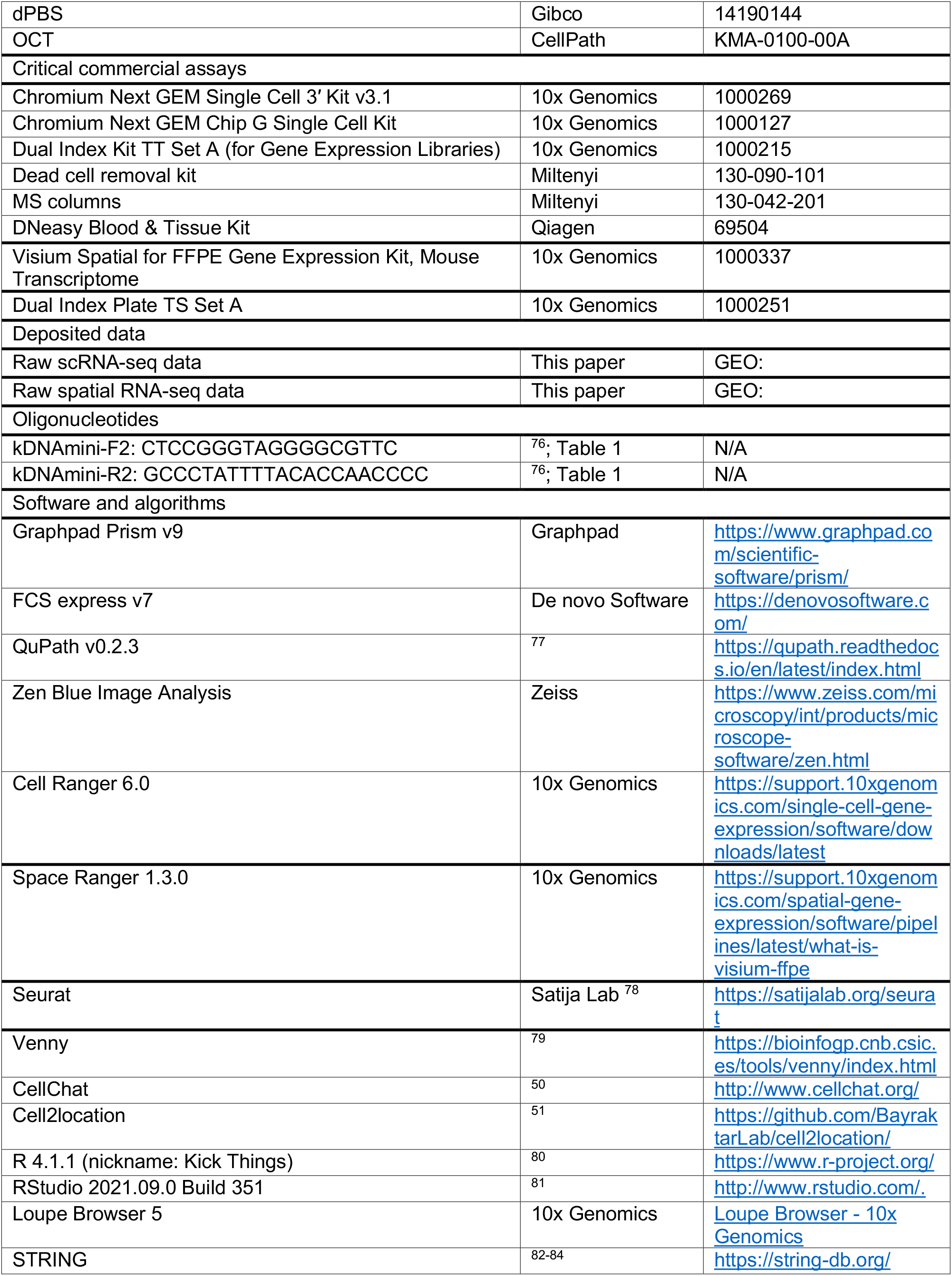

## RESOURCE AVAILABILITY

### Lead Contact

Further information, requests for resources and reagents should be directed to the lead contact, Paul M Kaye

### Materials Availability

RE9H luciferase-expressing *L. donovani* parasites generated in this study are available from the Lead Contact or Elmarie Myburgh with a completed Materials Transfer Agreement.

### Data and Code Availability

The sequencing data reported in this paper have been deposited in the GEO database under the accession code GEO: TBC Code used for analysis (including that for generating Figs from scRNA seq data) in this study is available at https://github.com/jipsi/leish-UVB

## EXPERIMENTAL MODEL AND SUBJECT DETAILS

### Mice

8–12-week-old, female C57BL/6J mice (RRID: IMSR_JAX_000664), originally obtained from the Jackson Laboratory were bred in house and maintained under pathogen free conditions at the Biological Services Facility at the University of York in accordance with the UK Home Office guidelines under the Animals (Scientific Procedures) Act 1986 (project licence No. P49487014). Results are reported in accordance with ARRIVE guidelines (**Supplementary Table S3**).

### Parasites and Infections

*L. donovani* (LV9; MHOM/ET/67/HU3) was originally isolated from a patient in Ethiopia in 1967 ^85^ and maintained by passage in B6.CD45.1.*Rag2*^−/−^ mice. WT or luciferase-expressing *L. donovani* parasites were used throughout this study where indicated. RE9H luciferase-expressing *L. donovani* were generated by transfecting log-stage promastigotes with linearized pRP-VH plasmid ^86^ containing a red shifted luciferase gene (Ppy-RE9H)^87^ and selecting with 50 μg mL−1 G418 (InvivoGen) as described previously ^88^. Passage mice were euthanised and the spleen was excised into 5mL Roswell Park Memorial Institute medium 1640 (Gibco; Life Technologies; 11875085) supplemented with 100μg/mL penicillin-streptomycin (Gibco; Life Technologies; 10378016) under aseptic conditions. The spleen was then homogenised using a glass tissue grinder and the cell suspension was centrifuged in a Heraeus Multifuge 3SR Plus (Thermo Scientific Heraeus®; 75004371) at 800rpm at 37°C for 5 minutes. The supernatant was transferred to a new tube and the pellet was discarded. The supernatant was then centrifuged at 3100rpm at 37°C for 10 minutes. The supernatant was discarded and the pellet was resuspended in 1mL of sterile ACK Lysing Buffer (Lonza; 10-548E) and incubated at 25°C for 5 minutes. 49mL of sterile RPMI/PS was then added and parasites were then centrifuged at 3100rpm at 37°C for 10 minutes. This wash with sterile RPMI/PS was completed three times. After the final wash, the supernatant was discarded and the pellet was resuspended in 1mL of sterile RPMI/PS. The parasite suspension was then taken up through a BD Microlance 3 needle (25G × 5/18” (BD; 300600) on a 1mL syringe (BD Plastipak; 303172) and dispensed 10 times to remove clumps. Amastigotes were then counted on a Thoma Counter (depth 0.2mm 1/400mm^2^, Weber England) and the following formula: (raw count/16) × 2×10^7^ was used to determine the concentration of the parasite suspension (parasites/mL). All mice were infected with a standard dose of 3×10^7^ parasites in 100μL i.v. via the lateral tail vein. Cage positions were shuffled daily within experimental racks.

## METHODS DETAILS

### Cell preparation and flow cytometry: Skin

Mice were shaved using a WELLA CONTURA HS61 Hair Clipper (Wella; HS61). Mouse flank skin (25mm × 25mm piece) was then collected into 5mL of 3% FBS. Subcutis layer (sub-cutaneous fat layer) removed by scraping with a rounded-edge scalpel (Swann Morton; 21-ref0507 and 11-ref0503) and rinsed with 3% FCS. Skin then minced into small pieces and transferred to a 5mL solution containing 3mg/mL Collagenase IV (Gibco; 17104-019) and 0.1mg/mL Dnase I (PanReac AppliChem, P120907) in RPMI1640 (Gibco; 11875085) + 10% FBS (HyClone™ Fetal Bovine Serum, South American Origin; GE healthcare Life Sciences; SV30160.03). Skin tissue was incubated for 2 hours at 37°C and then passed through a 40μM cell strainer and washed with 20mL of RPMI/PS. Skin single cell suspension was then centrifuged at 1700rpm for 10 minutes at 4°C, supernatant discarded and cell pellet resuspended in 200μL of MACS buffer, filtered through a 40μM cell strainer (Greiner bio-one; 542040) and transferred to a 96 U-well bottom plate (Sarstedt; 82.1582.001) for flow cytometric analysis or processed for scRNA-Seq.

### Cell preparation and flow cytometry: Spleen

For each mouse, the spleen was excised, weight was recorded in grams and collected into 10mL of 3% FBS in PBS. The spleen was dissociated through a 100μM cell strainer (Greiner bio-one; 542000) using the plunger end of a 5mL syringe (BD; 307731) and collected into a petri dish. Splenocyte single cell suspension was then transferred to a 15mL tube using a Pasteur pipette (SLS; 325685). Single cell suspension was then centrifuged at 1300rpm for 6 minutes at 4°C. Supernatants were discarded and cell pellet was resuspended in 1mL of ACK Lysis buffer (Lonza; 10-548E) and incubated at room temperature for 7 minutes. Samples were then washed with 9mL of 3% FBS and centrifuged at 1300rpm for 6 minutes at 4°C. Supernatants were discarded and cell pellet resuspended in 10mL of MACS buffer. 200μL of this single cell suspension was then transferred to a 96 U-well bottom plate (Sarstedt; 82.1582.001) for flow cytometric analysis.

### Flow cytometric staining

Samples were first blocked with Fc block for 10 minutes at 4°C incubated. Samples were then washed with MACS buffer (5mM EDTA, 0.01% FCS in PBS) and subsequently incubated with primary antibodies for 30 minutes at 37°C. Samples were then washed twice with MACS buffer and then acquired on a CytoFlex LX 375 (Beckman-Coulter) flow cytometer and FCS files were analysed in FCS express 7 Research (DeNovo Software).

### In vivo imaging

Mice were injected with 15mg/kg of D-luciferin (Syd labs; Cat MB000102-R70170) i.p and whole body imaged at 5 minutes post injection using field of view D on an IVIS Illumina XRMS series III (Caliper Life Sciences, PerkinElmer, UK) for 3 minutes. Mice were subsequently euthanised with CO_2_, shaved and the skin was removed and placed with the hydrophobic face up and imaged 10 minutes post injection using field of view C for 3 minutes.

### Ultraviolet B (UVB) exposure

Mice were shaved 24 hours prior to baseline measurements and first UVB dose. A UVB narrowband lamp (DermaHealer® compact; UAB Favoriteplus; Lithuania) was used to administer UVB treatment. The UVB lamp was calibrated using a UV-AB light meter (Extech-instruments; Model UV505). At 9cm away the UV-AB light meter measured UVB intensity to be 0.35mW/cm^2^ = 3.5J/s/m^2^. Therefore, a calibrated dose (at 9cm) of 500 J/m^2^ was administered to each mouse 3 times a week for 3 weeks prior to infection and 3 times a week for 2 weeks after infection ^31^.

### Skin assessment

Skin melanin and erythema were measured using a Mexameter MX-18 probe (EnviroDerm Services; UK) before UVB treatment and 24 hours post each UVB treatment session for the duration of the study. Animals were assessed in random order. The Mexameter MX-18 probe is equipped with LED light sources and a silicon diode detector for detecting reflected light from the skin. The Mexameter MX-18 measures the intensity of reflected green (568nm), red (660nm) and infrared (880nm). Melanin and erythema values are shown in one second as index numbers between 0 and 999. The Mexameter MX18 automatically calculates this as follows ^89^.

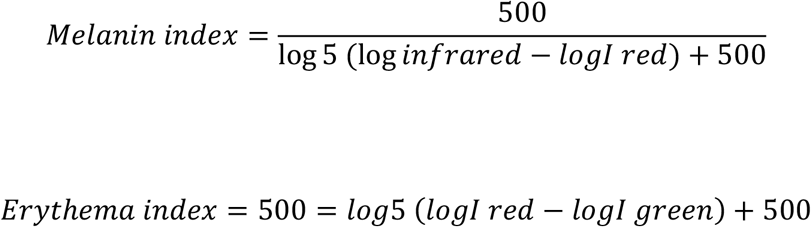

### gDNA extractions

6mm skin punch biopsies were collected and stored at −80°C until required. gDNA extractions were performed with the DNeasy Blood & Tissue Kit (Qiagen; 69504) as per manufacturer’s instructions. Briefly, skin punch biopsies were equilibrated to room temperature and then minced prior to the addition of 180μL of Buffer ATL and 20μL Proteinase K (per sample). Skin samples were then vortexed and incubated at 56°C overnight (or until the tissue was completely lysed). Washes and elution steps were performed as per manufacturer’s instructions. Quality of eluted gDNA was validated on a Thermo Scientific NanoDrop™ 1000 Spectrophotometer (ThermoFisher Scientific).

### qPCR

*Leishmania*-specific kinetoplastid DNA primers used in this study were previously characterized by ^76^ (Accession number AF103738) and used at a final concentration of 200 nM. 2 ng of gDNA were used per reaction (20μL reaction volume). Fast SYBR Green Master Mix (Applied Biosystems; 4385612) was used as per manufacturer’s instructions. Reactions were run on a QuantStudio 3 system; 96 well, 0.1mL (ThermoFisher Scientific) with a thermal cycle of 95°C for 20s, a cycling stage of 40 cycles of 95°C for 3s, 60°C for 30s, 95°C for 1s, 60°C for 20s, 95°C for 1s (data read at final step), followed by the standard melt curve stage. Data was analysed by ThermoFisher Connect cloud analysis software (ThermoFisher Scientific).

### FFPE

8mm biopsy punches of skin were collected into 4% PFA and stored at 4°C for 8 hours and then paraffin embedded in histosette I tissue processing/embedding casettes (Simport; M490-5) on the Leica ASP300S Fully Enclosed Tissue Processor (Leica Biosystems) and embedded on the Leica EG1150 H Modular Tissue Embedding Center (Leica Biosystems).

### H&E

Blocks were chilled prior to sectioning. 7μM sections were cut on a Leica Wax Microtome and placed into a water bath set to 45°C for 15 seconds. Sections were then collected onto Superfrost slides (ThermoScientific; J1800AMNZ) and allowed to dry overnight at RT. Slides heat fixed at 60°C for 2 hours in a sterilising oven (Leader Engineering; GP/30/SS/250/HYD, 08H028). Slides were allowed to cool down and then deparifinised with Histoclear II (SLS; NAT1334) for 5 minutes. Slides were equilibrated in 95% Ethanol for 3 minutes, 70% Ethanol for 3 minutes and distilled water for 3 minutes. Slides were then stained in Harris Haematoxylin (ThermoScientific; 6765001) for 3 minutes and then rinsed in tepid water for 5 minutes. Slides were dipped once in 1% acid-alcohol (HCl-EtOH; Sigma; 30721-2.5L-M; Fisher Scientific; E/0650DF/C17) and then equilibrated in distilled water for 3 minutes. Slides were then stained with 1% Eosin (Sigma-Aldrich; E4382-25G) for 3 minutes and then dipped in 50% Εthanol 10 times. Slides were then equilibrated in 70% Ethanol for 3 minutes, 95% Ethanol for 3 minutes and 100% Ethanol for 3 minutes. Slides were then cleared in Histoclear II (SLS; NAT1334) for 9 minutes. Slides were then mounted with Dibutylphthalate Polystyrene Xylene (DPX; Sigma-Aldrich; 06522-500ML) and coverslipped with 22 × 50 mm cover slips (SLS; MIC3226). Slides were dried overnight before scanned on the Zeiss Axioscan Z1 (Zeiss).

### Single cell isolation for scRNA-Seq

Skin tissue processed as above and dead cells removed using the dead cell removal kit (Miltenyi; 130-090-101) as per manufacturer’s instructions. An aliquot was taken and stained with CD45 and live/dead antibodies to check viability prior to library preparation.

### Single cell library preparation

Eluted cells after dead cell removal were washed in dPBS containing 0.04% BSA and resuspended at a concentration of approximately 1000 cells/μL. Library preparation was then performed following the Chromium Next GEM Single Cell 3’ Kit v 3.1 (10X Genomics; following the CG000315 Rev A user guide), where 10,000 cells are targeted for capture. Each library was sequenced on the Illumina NovaSeq 6000 platform, to achieve a minimum of approximately 20,000 reads sets per cell.

### Visium whole transcriptome spatial transcriptomics and processing

FFPE sections from non-infected -UVB, L. donovani infected -UVB and *L. donovani* infected +UVB mice (3 per group; total n=9) were cut onto 10X Genomics Visium slides and processed according to the Visium Spatial Gene Expression Reagent Kits for FFPE recommended protocol v1 (10X Genomics). Briefly, slides were stained with hematoxylin and eosin, imaged, and de-crosslinked. Mouse probes were added overnight and then extended and released. Libraries were prepared according to the manufacturer’s instructions and sequenced using the NovaSeq 6000 platform. Raw fastq files were aligned to the mouse genome mm10 using spaceranger software (10x Genomics). Associated image files were aligned onto slide specific fiducials using Loupe browser software (10X Genomics). Tissue regions were manually selected and a tissue x,y co-ordinate json file was created. Json files and image files were provided as input to the spaceranger count() function to generate counts and align them to spatial spots. Raw counts were normalised and analysed further. Gene expression enrichment in spatial spots were calculated using the AddModuleScore() function in Seurat and score greater than 0 was considered a positive score.

### Processing and quality control of scRNA-Seq data

FASTQ files were aligned using 10x Genomics Cell Ranger 6.1.0. Each library was aligned to an indexed mm10 genome using Cell Ranger Count. Generated .h5 files were loaded as Seurat objects ^78^ and quality controlled by removing cells with more than 10% mitochondrial reads. Cells were visualised as a scatter plot to visualise feature-feature relationships and subset accordingly. Counts for *Gm42418* and *AY036118* were removed from the samples these transcripts may represent library amplification noise ^90^. Next, each sample was regressed for ribosomal genes, mitochondrial reads, total RNA count, unique feature count using SCTransform() in Seurat using Gamma-Poisson generalised linear models, specifically, the method glmGamPoi (R package) ^91^. Finally, anchors for all samples were calculated using SelectIntegrationFeatures() and FindIntegrationAnchors() to integrate all groups into one integrated object.

### Dimensionality reduction and clustering analysis of scRNA-Seq data

First, principal components analysis was done to reduce the data into fewer dimensions. The top 15 principal components that explained most of the variance in the data were selected for downstream analysis. Low resolution Louvain clustering was used at a resolution of 0.4 to identify 16 clusters that were visualised using scatter plots either on principal component 1 and 2 or in UMAP space. Macrophage, T_NK, endothelial and LEC/LV cells were subset and re-clustered by repeating the above procedure to find further granularity in cellular phenotype. Comparison between cell types and groups were carried out by calculating differentially expressed genes using Wilcoxon-Rank sum test as employed in the FindAllMarkers() or FindMarkers() function in Seurat. Genes with p-values higher than 0.05 were discarded.

### Ligand-receptor interaction

Interactions between cell types were calculated using the R package Cellchat ^50^ that assigns a probability to each interaction and then conducts a permutation test. The interactions are based on a database of curated known interactions. We used a subset of interactions, namely, those falling under “secreted signalling” and “cell-to-cell contact” to model interaction network with a total of 1,073 interactions available to review on http://www.cellchat.org/. Finally, the L-R interactions were modelled based on law of mass action. Only significant interactions were presented.

### Spatial cell type de-convolution and graph-based clustering

Cell2location was used to de-convolute cell type per Visium spatial spot using the hyperparameters N_cells_per_location=30, detection_alpha=20 using our own single cell RNA seq data set (this paper). First, the reference cell type signatures were estimated using a negative binomial regression model by training the model for 1,000 epochs. The reference signature model was then used to deconvolute the spatial data with abovementioned hyperparameters by training cell2location model for 30,000 epochs. Finally the 5% quantile of the posterior distribution wherein the model has high confidence was used to infer the value of cell abundance per spatial spot. Next, the abundances were subset for a selection of cells that were most abundant by removing all cell types whose max value was lower than the median of all the max values calculated per cell type. The abundances were then used to generate a correlation matrix which was then used as a distance matrix (1-corr) to create a graph. A minimum spanning tree was calculated to obtain the shortest distance between the graph and clusters were calculated using edge.betweenness.

### STRING and GSEA

Differentially expressed gene names were submitted to StringDB (https://string-db.org/). The full STRING network (the edges indicate both functional and physical protein associations) was selected for the analysis. All default interaction sources with a medium confidence interaction score of 0.4 were selected (text-mining was excluded from the analysis). Gene set enrichment analysis (GSEA) was conducted using HALLMARK gene lists (https://www.gsea-msigdb.org/gsea/index.jsp) and the calculated FDR q-value was converted to −LOG_10_(FDR) for presentation purposes.

## QUANTIFICATION AND STATISTICAL ANALYSIS

Animals were reandomised to treatment groups. Downstream data analysis was unblinded and conducted using quantitative methodologies as described above. No animals or data points were excluded from the analysis. Statistical analyses were performed in Prism 9 (v9.01; GraphPad Software). Data were tested for normality using the D-Agostino & Pearson method. Data are presented as the mean ± SEM or the median ± IQR as indicated. Power calculations (type I/II error rate, where α = 0.05 and power = 80%) determined the sample size to include 5 mice per group. Sample sizes in each plot has been listed in the Fig. Legends where appropriate. P values are shown as *p < 0.05, **p < 0.01, ***p < 0.001 and ****p < 0.0001. Groups with two or more dependent variables were compared using a One-Way ANOVA with Tukey’s multiple comparisons test or a Two-Way ANOVA with Sidak’s multiple comparisons test.

