## Supplementary Figures 1-9 for "UVB modifies skin immune-stroma cross-talk and promotes effector T cell recruitment during cryptic *Leishmania donovani* infection"

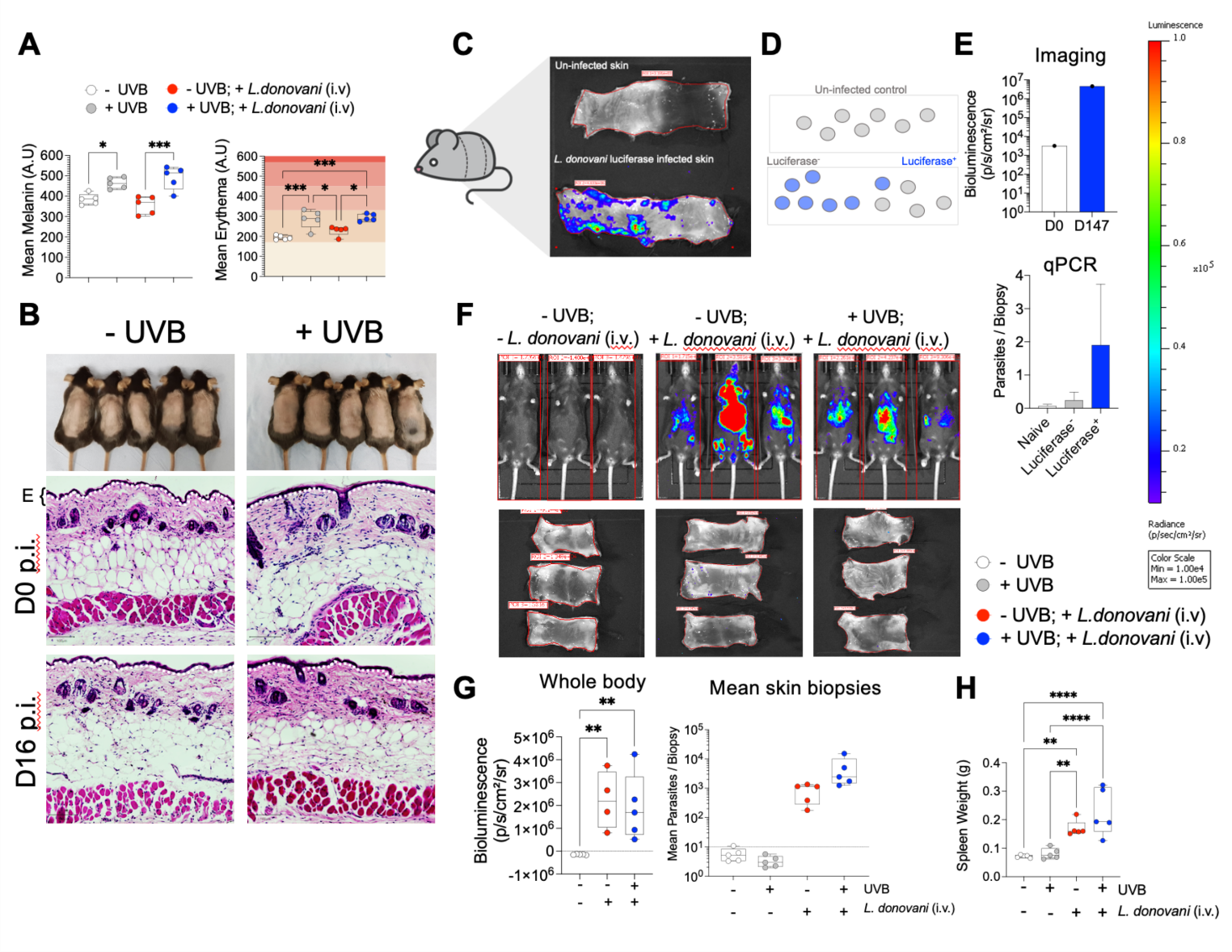

**Fig. S1: Low UVB dose elicits skin histological changes during *L. donovani* infection and PCR detects parasites in patches associated with *in vivo* bioluminescence**

**A)** C57BL/6J females shaved and ear notched 24 hours prior to baseline measurements and first UVB dose (n = 5 mice per group). Average melanin and erythema measured in arbitrary units (A.U) **B)** Photos and 7μM H&E skin sections from unexposed and UVB-exposed groups 16 days post infection (p.i.) and 36 days post first UVB dose. ‘E’ denotes epidermis along with white dotted line. Scale bars on H&E images: 100μM. **C)** B6.CD45.1Rag2^-/-^ female infected with 3x10^7^ *L. donovani* luciferase amastigotes i.v.; D0 (n=1), D147 (n=1). Mice were injected with D- luciferin and skin imaged ~ 8 mins post injection; FOV: C for skin; exposure: 3 mins smoothing: 7x7, all images adjusted to the same colour scale of bioluminescence - min: 1x10^4^ max: 1x10^5^; ROI: free-form shaped. **D)** 8mm skin biopsy punches collected from each mouse. 7 biopsies from the uninfected control, 7 biopsies from luciferase^+^ (blue) and 4 biopsies from luciferase^-^ (grey) areas. **E)** Bioluminescence in skin (left) and qPCR used to determine parasite burdens in individual skin biopsies (right). **F)** C57BL/6J females shaved and ear notched 24 hours prior to baseline measurements and first UVB dose (n = 4-5 mice per group). Mice exposed with UVB-light at 500J/m^2^ three times weekly for 3 weeks. Mice were infected with 3x10^7^ *L. donovani* luciferase amastigotes i.v. and exposed or not with UVB-light at 500J/m^2^ three times weekly for remaining 2 weeks. Mice were injected with 15mg/kg of D- luciferin i.p. and whole-body bioluminescence measured 5 mins post injection. **G)** 4mm skin biopsy punches were collected from each mouse and 2ng gDNA used as input for qPCR.. **H)** Spleen weights on day 16 p.i.. Boxplots show the minimum, the maximum, the sample median, and the first and third quartiles. One-way ANOVA with Tukey’s multiple comparisons test; *p < 0.05, ** p < 0.01, ***p < 0.001, ****p < 0.0001. Data shown are representative of two independent experiments.

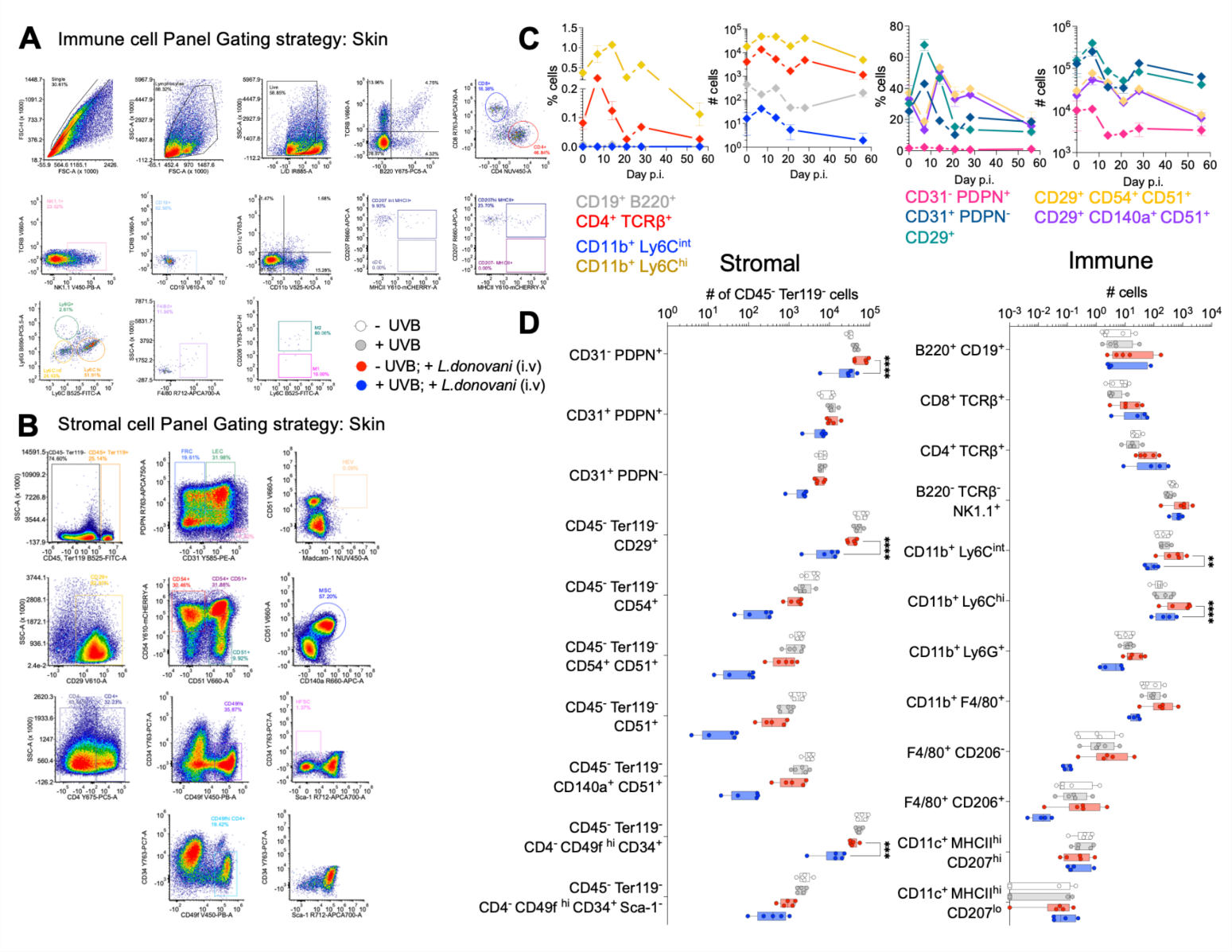

**Fig. S2: Kinetics and abundance of immune and stromal cell frequencies and abundance during *L. donovani* infection**

C57BL/6J females infected with 3x10^7^ *L. donovani* luciferase amastigotes i.v.; n=5 mice per timepoint, spleen and skin tissue processed for flow cytometry; error bars depict mean $\pm$SEM, spleen cell numbers adjusted to organ size; skin cell numbers adjusted to whole skin area (Collins *et al.,* 2016 estimate: 36cm^2^). **A)** Gating strategy for immune cell panel in the skin. **B)** Gating strategy for stromal cell panel in the skin. **C)** Frequency and cell numbers of immune and stromal cells measured in the skin by flow cytometry at day 0, 7, 14, 21, 28 and 56 p.i.. Data shown are from one experiment (A – C). Immune cells defined as:

- Patrolling monocytes: B220^-^ TCRβ^-^ CD11c^-^ CD11b^+^ Ly6C^int^ Ly6G^int^
- Inflammatory monocytes: B220^-^ TCRβ^-^ CD11c^-^ CD11b^+^ Ly6C^hi^
- CD4^+^ T cells: B220^-^ TCRβ^+^ CD4^+^
- B cells: B220^+^ TCRβ^-^ CD19^+^
- Stromal cells defined as:
- Skin stromal cells: CD45^-^ Ter119^-^ CD29^+^ CD31^-^ Pdpn (gp38)^+^
- Blood endothelial cells (BEC): CD45^-^ Ter119^-^ CD29^+^ CD31^+^ Pdpn (gp38)^-^
- Lymphoid endothelial cells (LEC): CD45^-^ Ter119^-^ CD29^+^ CD31^+^ Pdpn (gp38)^+^
- Activated endothelial cells (AEC): CD45^-^ Ter119^-^ CD29^+^ CD51^+^ CD54^+^
- Mesenchymal stromal cells (MSC): CD45^-^ Ter119^-^ CD29^+^ CD51^+^CD140a^+^

**D)** C57BL/6J females shaved and ear notched 24 hours prior to baseline measurements and first UVB dose (n = 4-5 mice per group). Mice exposed to UVB light at 500J/m^2^ three times weekly for 3 weeks. Mice were then infected with 3x10^7^ *L. donovani* amastigotes i.v. Mice exposed to UVB light at 500J/m^2^ three times weekly for remaining 2 weeks. Skin tissue collected for flow cytometric analysis. Plots show the numbers of stromal and immune cell populations in the skin at day 16 p.i. Box plots show the minimum, the maximum, the sample median, and the first and third quartiles. One-way ANOVA with Tukey’s multiple comparisons test; * p < 0.05, ** p < 0.01, ****p < 0.0001. Data shown are representative of two independent experiments (D).

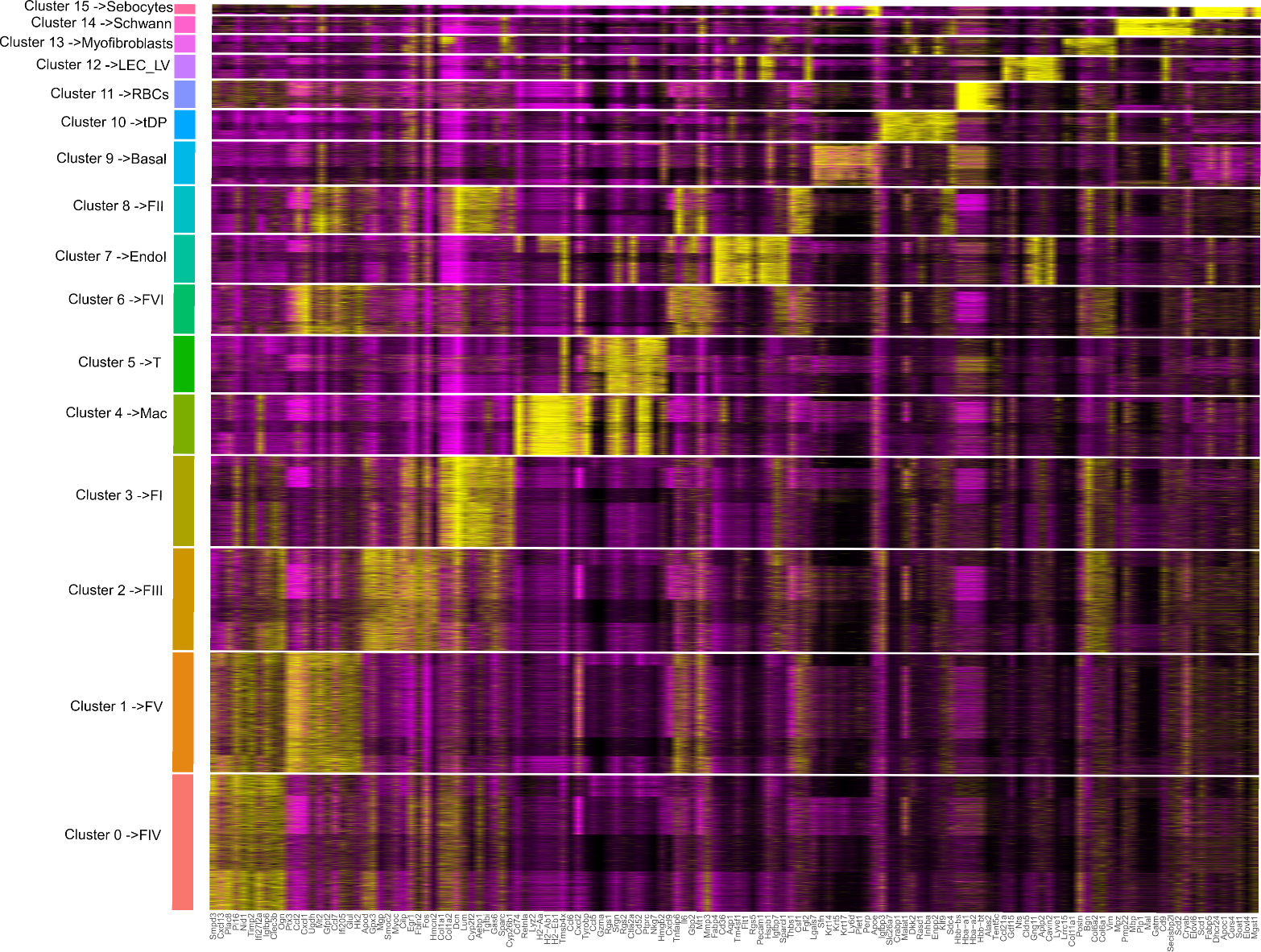

**Fig. S3: Top features of scRNA-seq clusters/cell type**

Heatmap showing top 10 genes per cluster for all 16 clusters/cell types

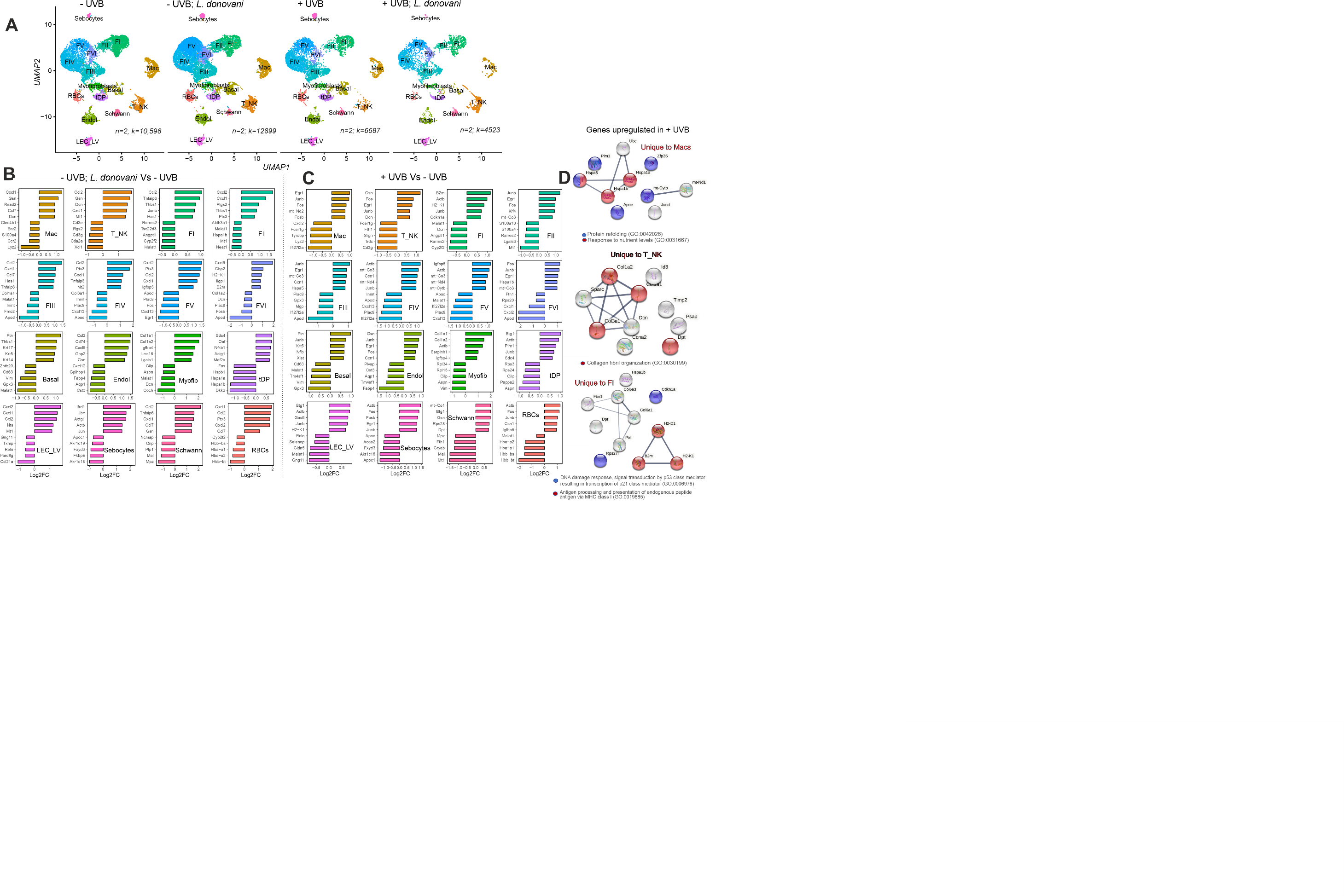

**Fig. S4: DEGs associated with *L. donovani* infection in -UVB mice and with UVB exposure alone**

**A)** UMAP plots split across all comparison groups coloured by cell type where n indicates number of mouse per group (pooled) and k number of cells per group. **B and C)**. Bar plots showing top 5 and bottom 5 genes regulated (expressed as log2fold change) in all cell populations for infected -UVB mice vs. uninfected -UVB mice (B) and for uninfected +UVB vs. -UVB mice (C). **D)** Network diagrams coloured by gene ontology terms of genes uniquely upregulated in macrophages, T cells and FI fibroblasts in +UVB vs. -UVB mice.

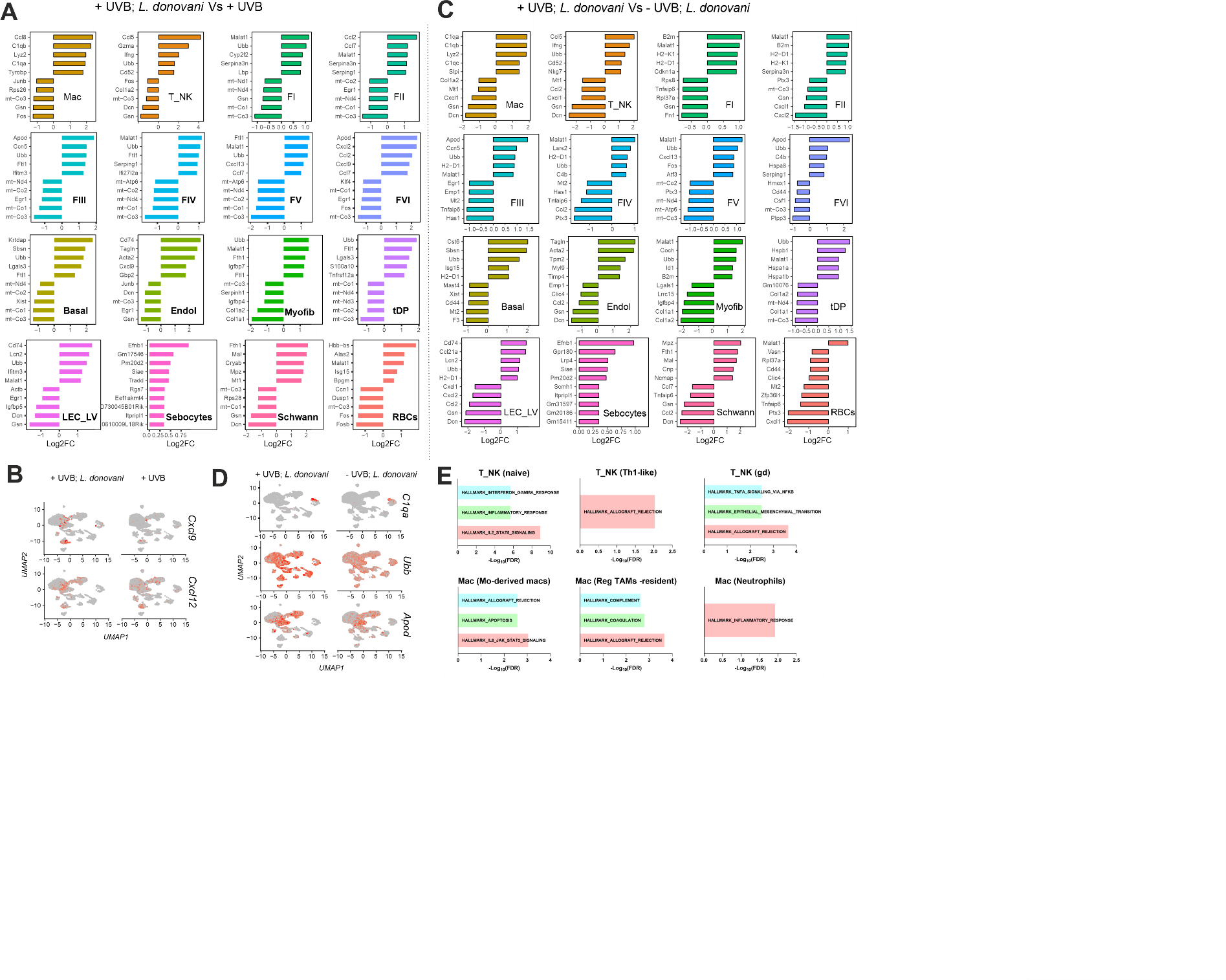

**Fig. S5: DEGs associated with *L. donovani* infection in UVB exposed mice**

**A)** Bar plots showing top 5 and bottom 5 genes regulated (expressed as log2fold change) in all cell populations for infected +UVB mice vs. uninfected +UVB mice. **B)** UMAP plots showing *Cxcl9* and *Cxcl2* expression between the above comparison. **C)** Same as A but for infected +UVB mice vs. infected -UVB mice **D)** UMAP plots showing *C1qa*, *Ubb* and *Apod* expression between the comparison in C. **E)** GSEA enrichment shown as bar plots depicting negative log FDR for genes upregulated in infected +UVB mice versus infected -UVB mice for T and macrophage sub-populations.

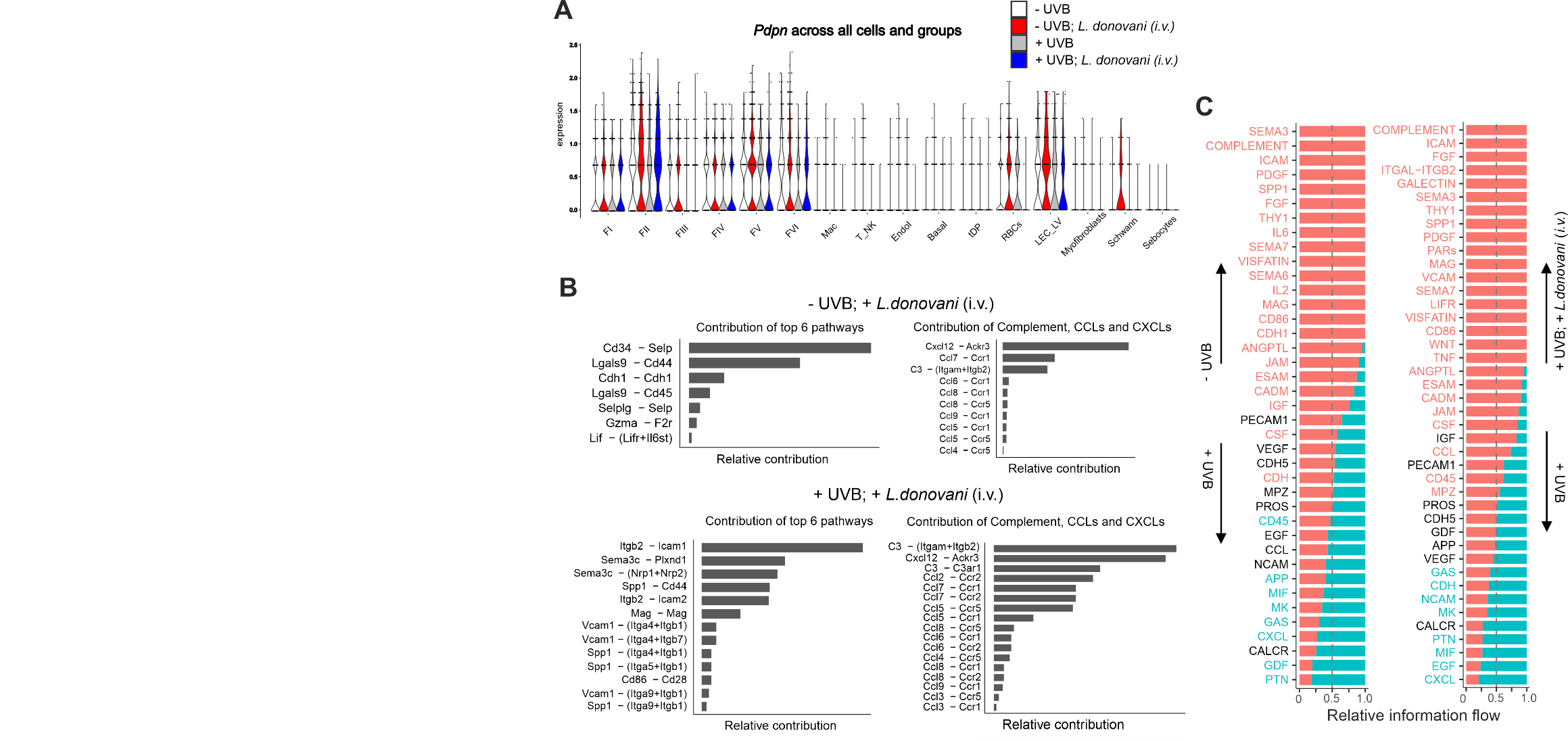

**Fig. S6: *Pdpn* transcripts expression across cell types and groups and receptor-ligand interactions in infected +UVB mice vs. infected -UVB mice**

**A)** Violin plot showing normalised gene expression of *Pdpn* across cell groups. **B)** Relative contribution of each ligand-receptor pair to the overall signalling networks uniquely up in infected -UVB mice vs. infected +UVB mice (left). Contributions of complement and *Ccl-Ccr* network (right). **C)** Bar plots showing pairwise comparison of information flow between inferred signalling networks with pathways in red more enriched in groups indicated with upward arrow and vice versa.

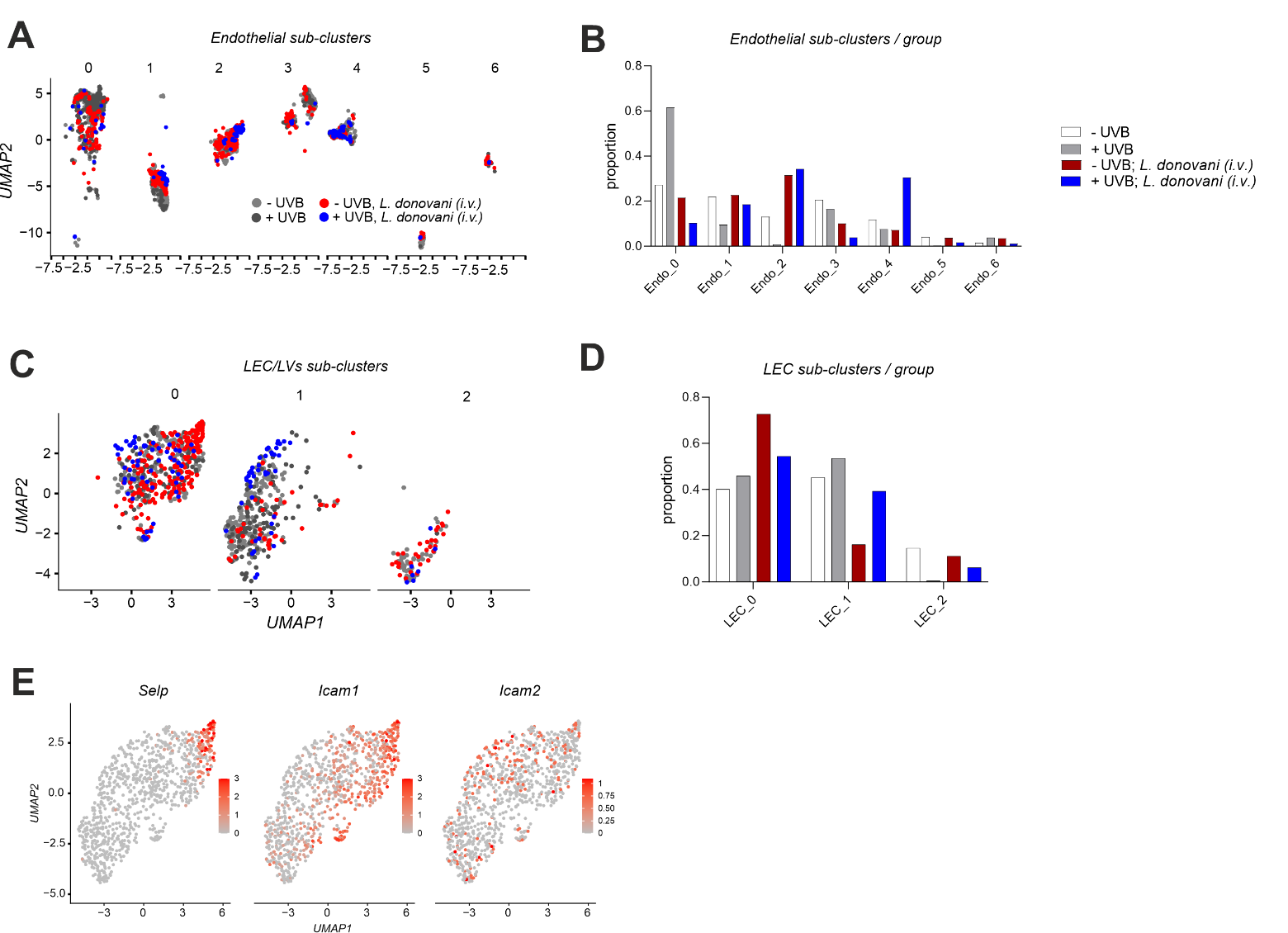

**Fig. S7: Endothelial and LEC/LV sub-clustering**

**A)** UMAP plots of endothelial cells indicating treatment group. **B)** Relative proportion of endothelial sub-clusters by treatment group. **C).** UMAP plots of LEC_LV indicating treatment group. **D)** Relative proportion of LEC_LV sub-clusters by treatment group. **E)** UMAP plot of sub-clustered LEC_LVs highlighting cells expressing *Selp*, *Icam1* and *Icam2.*

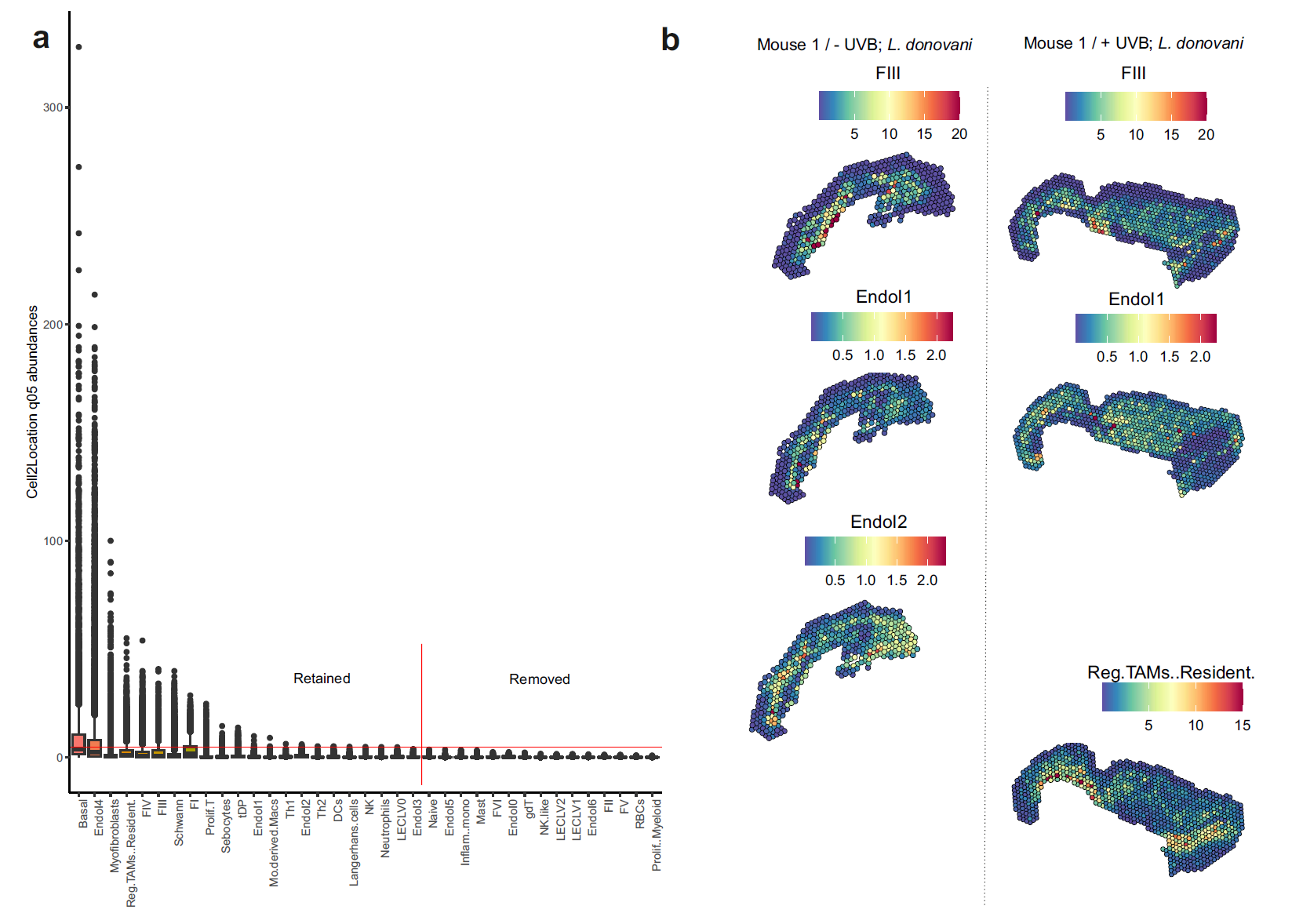

**Fig. S8: Spatial localisation of cell2location cell type abundances.**

**A)** Cell2location predicted abundances for all 10,527 visium spots (all groups combined) are plotted in the order of their max value. Finally, a horizontal red line is drawn at y=median(max(all_cell_types)). A second vertical red line is drawn to indicate that the cell types to the left were included in the downstream analysis.

**B)** Spatial maps of cell type abundances for FIII, Endo1, Endo2 for mouse 1 from the  *L. donovani* infected -UVB mice and FIII, Endo1, Reg TAM Resident macrophages for mouse 1 from the *L. donovani* infected +UVB mice.

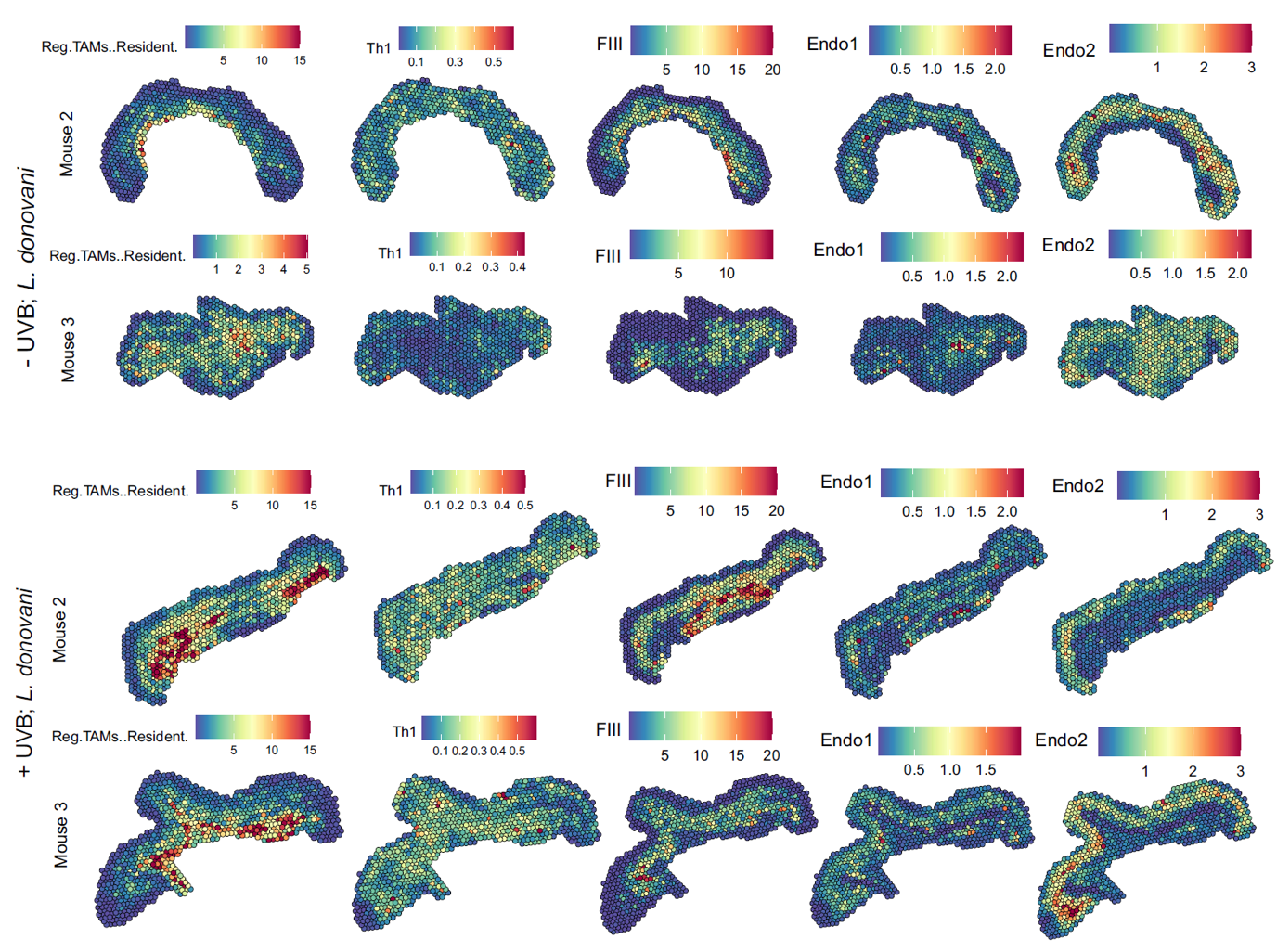

**Fig. S9 Spatial localisation of cell2location cell type abundances.**

Spatial maps of cell type abundances for Reg TAM Resident macrophages, Th1, FIII, Endo1, Endo2 cells for mouse 2 and 3 from *L. donovani* infected -UVB (top two rows) and *L. donovani* infected +UVB (bottom two rows) cohorts.
