## Supplementary Table 3 for "UVB modifies skin immune-stroma cross-talk and promotes effector T cell recruitment during cryptic *Leishmania donovani* infection"

Supplementary Table S3

ARRIVE Essential 10

| Study design | a) For each experiment; +UVB is compared to -UVB or +UVB +*L.donovani* is compared to -UVB+ *L.donovani* infected groups. -UVB groups are the control groups in both comparisons b) Experimental unit:   cage of 5 mice per group |
| --- | --- |
| Sample size | a) 5 mice per group, 20 mice in total per experiment. Total of 20 mice used in this study. b) Power calculations (type I/II error rate, where α = 0.05 and power = 80%) determined the sample size to include 5 mice per group. |
| Inclusion and exclusion criteria | a) Female mice were exclusively used in this study to avoid confounding skin inflammation that often results from male aggression. Day 16 timepoint post infection was chosen, due to hair regrowth occurring post day 16 and confounding Mexameter readings. b) No data points were excluded in this study c) Figure legends report exact *n* used in each analysis |
| Randomisation | a) Simple randomisation method was used to assign mice to control and treatment groups b) Confounders for UVB exposure was controlled for measuring the UVB dose prior to exposing mice to UVB light and alternating the order in which mice were exposed to UVB. Confounders for erythema and melanin readings were controlled for taking measurements of each group in a random order prior to UVB exposure. Cages were shuffled daily by BRF technical staff. |
| Blinding | Downstream data analysis was not blinded to treatment group |
| Outcome measures | Percentages of Immune and stromal cell phenotypes were quantified. Skin erythema and melanin content was quantified. |
| Statistical methods | a) Statistical methods for each analysis are described in Figure legend and methods of this study. b) Data passed the normality test using the D-Agostino & Pearson method. |
| Experimental animals | C57BL/6J female mice used in all experiments at 8 – 12 weeks old |
| Experimental procedures | All procedures are described in detail in the methods section on this study |
| Results | All data is reported in the results section of this study and listed for each analysis in the Figure Legends |
